# Comparative analyses of prolonged stress responses of breast carcinoma cells after hypofractionated irradiation *in vitro* and *in vivo*

**DOI:** 10.64898/2026.05.12.724504

**Authors:** Nirina Sivakumar, Gerhard Fritz

**Affiliations:** Institute of Toxicology, Medical Faculty and University Hospital Duesseldorf, Heinrich-Heine University Duesseldorf, Building. 22.21, Moorenstrasse 5, 40225 Duesseldorf, Germany

**Keywords:** Hypofractionated irradiation, radiation resistance, breast cancer, DNA damage response, DNA repair, cell death, inflammation, tumor microenvironment

## Abstract

The anticancer efficacy of radiotherapy (RT) is limited by acquired radioresistance (RR). Here, we aim to characterize prolonged responses of breast carcinoma cells to hypofractionated irradiation (hFI). To this end, murine mammary 4T1 tumor cells (4T1^WT^) were subjected to a clinically oriented hFI protocol (56 Gy cumulative dose) to select radioresistant cells *in vitro* (4T1^RS^). Furthermore, hFI of subcutaneously growing 4T1^CTR^ tumors (hFI; 24 Gy cumulative dose) was performed to radioselected 4T1^IR^ cells *in vivo*. Following single irradiation *in vitro*, radioselected 4T1^RS^ cells revealed increased proliferation, attenuated G2/M arrest and reduced apoptosis as compared to parental 4T1^WT^ cells. Moreover, 4T1^RS^ cells showed increased expression of DNA-damage response (DDR)-related proteins (pKAP1, pCHK2, γH2AX) and improved DSB repair efficiency as demonstrated by nuclear γH2AX foci analyses. The mRNA expression of factors regulating cell cycle progression, DDR, apoptosis and oxidative stress was substantially different between both cell variants *in vitro*. Ten days after hFI of *in vivo* growing tumors, residual DNA damage and apoptosis were increased in the radioselected 4T1^IR^ tumors, whereas proliferation was reduced as compared to non-irradiated 4T1^CTR^ control tumors. Both irradiated and non-irradiated tumors revealed complex differences in the mRNA expression profile of susceptibility- and metastasis related genes, including *GADD45a, DUSP1, CDKN1a* and *NQO1* as well as *CD44* and Rho-related factors, respectively. Moreover, hFI stimulated the infiltration of MPO-positive immune cells into tumor tissue while the presence of CD3-positive cells was reduced in the tumor area. In addition, hFI *in vivo* resulted in a dysregulated mRNA expression of various immune cell markers, Rho-regulatory factors, tissue remodeling molecules and cell adhesion factors. Summarizing, we identified long-lasting adaptive changes following hFI *in vitro* and *in vivo* that are associated with DNA replication, DNA repair, senescence and apoptosis as well as immune cell infiltration and tissue remodeling.

## Introduction

Radiotherapy (RT) plays a pivotal role in the treatment of malignant diseases by selectively inducing cytotoxic effects in tumor tissue while largely sparing exposure of normal healthy tissue. Usually, RT is performed as fractionated irradiation (FI), using a daily dose of 2 Gy and a cumulative dose of approximately 50-70 Gy ^1, 2^. Hypofractionated irradiation (hFI) regimen include a lower number of treatment cycles, yet with single doses of 4-6 Gy per fraction ^3^ being applied. hFI increases the likelihood that cycling tumor cells are exposed in particularly vulnerable phases of the cell cycle (i.e., S/G2), thus enhancing the therapeutic efficacy of RT. In addition, hFI allows a more efficient repair of sublethal damage in normal tissue ^4^, thus reducing the severeness of adverse radiation-induced normal tissue injury.

Ionizing radiation (IR) can damage various cellular macromolecules. However, the genomic DNA is the primary (i.e., most relevant) target structure in the context of RT-based anticancer therapy ^5, 6^. IR induces various types of DNA damage, including oxidative base modifications, DNA single-strand breaks (SSBs) and DNA double-strand breaks (DSBs) ^7^. DSBs are highly cytotoxic lesions ^8^ and potent activators of the DNA damage response (DDR), which regulates the balance between survival- and cell death-related pathways by activating cell cycle checkpoints as well as DNA repair- and/or apoptosis-related mechanisms ^9–13^. The PI3 kinase-like protein kinases Ataxia telangiectasia mutated (ATM) and ATM and Rad3-related (ATR) are the key regulators of the DDR ^12, 14^. ATM is recruited to DSB by help of the MRN complex (Mre11-Rad50-Nbs) ^15^. ATR is activated particularly as a result of DNA replication blockage caused e.g. by DNA crosslinks or DSBs ^16, 17^ and is recruited to the arrested replication fork ^18^. Targets of ATM and ATR include checkpoint kinases 1 and 2 (CHK1/2), which regulate the activation of G1/S, intra-S, and G2/M cell cycle checkpoints. Hence, CHK1/2 are considered as promising targets for novel anticancer therapeutics ^19^. Moreover, ATM/ATR-regulated signaling also affects the function of the tumor suppressor p53, which defines the balance between survival, senescence and cell death ^11, 20^. Repair of DSB occurs through homologous recombination (HR), which is a high fidelity repair mechanism occurring in S- and G2-phase, or non-homologous end joining (NHEJ), which is more error-prone ^8^ and present in all phases of the cell cycle, including G0/G1 ^9^. In the case of defective NHEJ, alternative end joining pathway can complement DSB repair ^21, 22^. HR requires BRCA2 (Breast Cancer Associated gene 2) ^23, 24^, which is particularly relevant as a tumor suppressor in hereditary forms of breast cancer ^25^. Lack of BRCA2 increases the sensitivity of tumors to PARP inhibitors (concept of synthetic lethality) ^26, 27^. Considering the multiple biological effects of radiation, molecular mechanisms contributing to acquired radiation resistance (RR) are rather diverse ^28–31^. Thus, RR of malignant cells has been associated with mechanisms of cell cycle progression ^32, 33^, p53- and apoptosis-associated factors ^34–39^, PI3K/AKT/mTOR signaling ^40–42^, NF-κB-^43^, HIF1-^44^ and EGFR-associated signaling pathways ^45^, cell adhesion mechanisms ^46^, NRF2-regulated anti-oxidative stress responses ^47^, and, in particular, mechanisms of the DDR and DNA repair ^48–52^. In addition, the tumor immune microenvironment (TME) influences signaling pathways promoting acquired RR by immunosuppressive features ^53^. Moreover, loss of p53 plays a pivotal role in promoting RR of malignant cells due to G1/S checkpoint failure ^54^. However, it is unclear, to which extent these different molecular mechanisms are clinically relevant as radioresistance factors.

Most important, the majority of nowadays available preclinical RR-related data was obtained from single irradiation experiments, whereas repeated FI/hFI is used under clinically relevant therapeutic situation. Aiming to identify clinically relevant hFI-related resistance factors, it is rational to assume that the specific experimental conditions used to select radioresistant tumor cells in preclinical *in vitro* and/or *in vivo* models significantly influences the spectrum of identifiable RR-related susceptibility factors. Moreover, having in mind that the TME is known to impact radiation sensitivity ^55–58^ it is feasible that IR-induced stress responses of tumor cells *in vitro* substantially differ from their responses observed *in vivo*. Therefore, in our present study we aimed to systematically compare hFI-induced stress responses *in vitro* and *in vivo*. To this end, we employed murine 4T1 mammary carcinoma cells for hFI *in vitro*, aiming to identify long-lasting alterations influences their IR responsiveness to a single IR treatment. In addition, we employed a syngeneic mouse model of subcutaneously growing 4T1 tumors (xenograft model) to identify prolonged tumor-associated alterations caused by hFI of the locally growing tumors *in vivo*.

## Materials and Methods

### Cell culture conditions and radiation treatment

Murine 4T1 mammary carcinoma cells originate from the American Type Culture Collection (ATCC) (Manassas, VA, USA) and were cultured in RPMI-1640 medium (Sigma, Steinheim, Germany) containing 10 % heat-inactivated fetal bovine serum (Biochrom, Berlin, Germany) and 1 % penicillin/streptomycin (Sigma, Steinheim, Germany) at 37 °C in a humidified atmosphere containing 5 % CO_2_. For radioselection, 4T1 cells were exposed to a hypofractionated irradiation (hFI) scheme over 5 weeks (4 Gy per single ionizing radiation (IR) exposure; 1-3 IRs per week; cumulative radiation dose of 56 Gy) **(see Fig. 1A)**. For comparative analysis of the response of the non-irradiated wildtype and radio-selected cells, they were seeded in dishes and grown to optical confluency for 48 h. Afterwards cells were irradiated with 4-8 Gy using the ^137^Cs source Gammacell 3000 (Nordion, Ottawa, ON, Canada).

**Figure 1:**
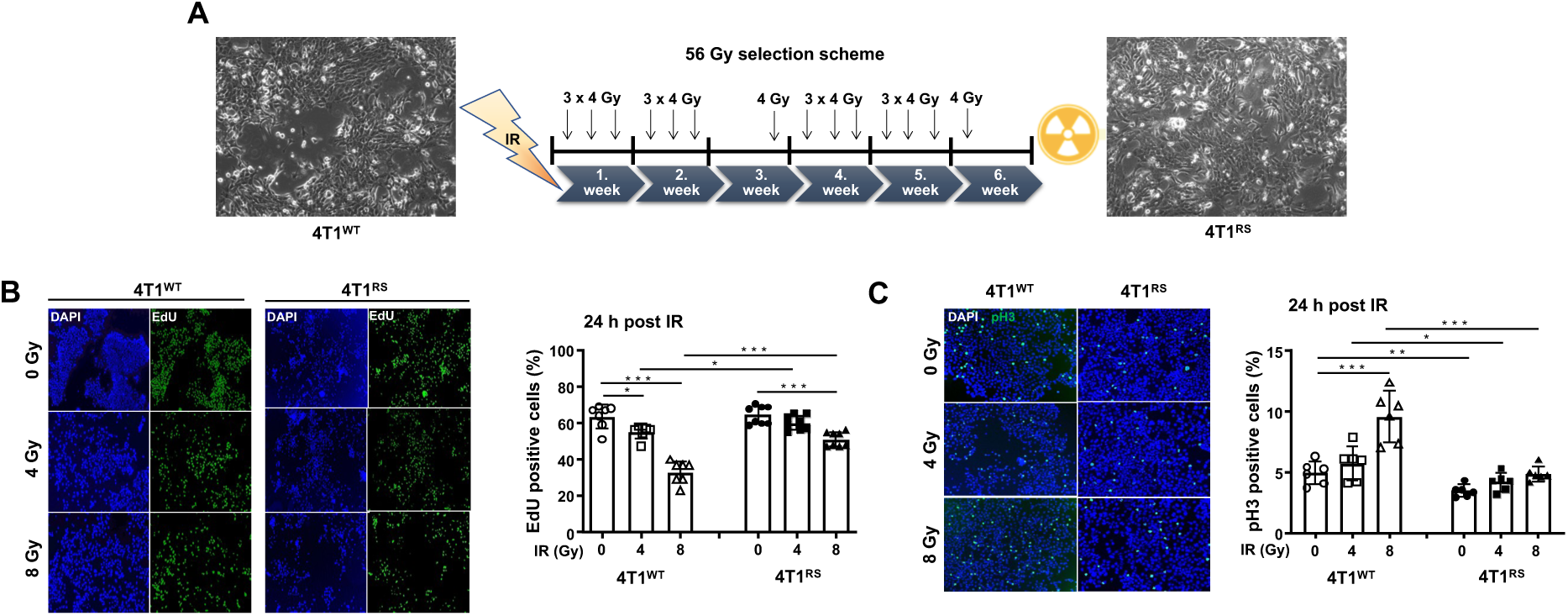
Comparative analysis of proliferation following irradiation treatment of 4T1^WT^ versus 4T1^RS^ *in vitro*. **A**: Schematical illustration of clinically oriented *in vitro* hypofractionated irradiation scheme used. Parental murine 4T1 breast cancer cells were exposed to hFI over 5 weeks (4 Gy per treatment; 1-3 irradiations/week; cumulative dose of 56 Gy). **B:** Logarithmically growing 4T1 and radioselected 4T1^RS^ cells were irradiated with 4-8 Gy. After a post-incubation period of 24 h cell proliferation was assayed by monitoring EdU incorporation. Representative pictures are shown on the left. Quantitative data shown in the histogram on the right are the mean ± SD from n=4 independent experiments each performed in biological duplicates (N=7-8). Statistical analysis was performed using unpaired, two-tailed Student’s t-test. *, p ≤ 0.05; **, p ≤ 0.01; ***, p ≤ 0.001 (irradiated cells as compared to the corresponding non-irradiation control). **C:** Logarithmically growing 4T1^WT^ and radioselected 4T1^RS^ cells were irradiated with 4-8 Gy. After a post-incubation period of 24 h mitotic index was determined as described in methods. Representative pictures are shown in the left. Quantitative data are shown as the mean ± SD from n=3 independent experiments each performed in biological duplicates. Statistical analysis was performed using unpaired, two-tailed Student’s t-test. *, p ≤ 0.05; **, p ≤ 0.01; ***, p ≤ 0.01 (irradiated cells as compared to the corresponding non-irradiated control).

### Flow cytometry-based analysis of cell cycle progression and cell death

For flow cytometry-based analysis of cell cycle distribution, cells were trypsinized, washed, and resuspended in PBS. Following fixation with ice-cold ethanol (80 %) (≥20 min, −20 ^°^C), cells were pelleted (800 x *g*, 10 min, 4 ^°^C) and suspended in PBS. DNase-free RNase A (Serva Electrophoresis GmbH, Heidelberg, Germany) was added (1 μg/μl, 1 h, RT) to degrade RNA. Afterwards, nuclear DNA was stained with propidium iodide (PI) (50 μg/ml, Sigma-Aldrich, St. Louis, MO, USA). Quantification of the percentage of cells presents in the SubG1-fraction (indicative of apoptotic cell fraction), G1-phase, S-phase, and G2/M-phase was performed using BD Accuri™C6 Flow Cytometer (BD Biosciences, San Jose, CA, USA).

### Analysis of cell proliferation - EdU incorporation and mitotic index

To analyze S-phase activity, the incorporation of the thymidine analogue 5-ethynyl-2’-deoxyuridine (EdU) into newly synthesized DNA was monitored. 24-72 h after IR, cells were pulse-labelled with EdU (1:500; 2 h) using the EdU-Click 488 Kit (Baseclick GmbH, Neuried, Germany) according to the manufacturer’s instructions. In addition, mitotic activity of cells was investigated by measuring the percentage of phospho-histone 3 (pH3 (Ser10)) positive cells. 24-72 h after IR treatment, cells were fixed with 4 % formaldehyde/PBS followed by incubation with ethanol (≥20 min, −20 °C). After blockage of unspecific antibody binding (5 % BSA in 0.3 % Triton X-100/PBS (1 h, RT)), anti-Ser10 pH3 antibody (1:1000, 4 °C, Abcam, Cambridge, MA, USA) was added (overnight incubation, RT). After washing, incubation with secondary Alexa Fluor^®^ 488 labeled goat anti-rabbit was performed (2 h, RT). After DNA counterstaining with DAPI-containing Vectashield^®^ (Vector Laboratories, Burlingame, CA, USA), the percentage of EdU-positive and pH3-positive cells was determined by Olympus BX43 microscope (Olympus, Hamburg, Germany) (10x objective). Data are shown as the mean ± SD of n=3-4 independent experiments each performed in biological duplicates with ≥100 nuclei being analyzed per experimental condition.

### Analysis of DNA damage formation and repair

The frequency of nuclear foci formed by S139 phosphorylated H2AX (γH2AX foci), which is a well-accepted surrogate marker of DNA double-strand breaks (DSBs)^59, 60^, was assayed by immunocytochemistry. Moreover, the appearance of nuclear 53BP1 foci, which is another marker of DSBs ^61, 62^ that are subject of DNA repair by non-homologous end joining (NHEJ), was additionally determined. 48 h after seeding onto coverslips, cells were irradiated and fixed at specific time points after irradiation (30 min, 2 h, 6 h, 24 h) with 4 % formaldehyde/PBS (15 min, RT), followed by incubation with ice-cold ethanol (≥20 min, −20 °C). After blocking (5 % BSA in PBS/0.3 % Triton X-100 (1 h, RT), co-staining with γH2AX antibody (Millipore) and 53BP1 antibody (Cell Signaling Technology, Cambridge, UK) was performed (1:500, overnight, 4 °C). After incubation with the secondary fluorescence-labeled antibodies Alexa Fluor^®^ 555 und Alexa Fluor^®^ 488 (1:500, 1 h, RT, in the dark), cells were mounted in Vectashield^®^ (Vector Laboratories, Burlingame, CA, USA) containing DAPI. The number of nuclear γH2AX and 53BP1 foci was scored microscopically (Olympus BX43 fluorescence microscope). Only nuclei with distinct foci were evaluated. γH2AX pan-stained nuclei, which are indicative of apoptotic cells, were excluded from the analyses. Furthermore, cells with micronuclei were counted. Data are shown as the mean ± SD of four independent experiments with ≥50 nuclei analyzed per experimental condition.

### Western Blot analyses

Total cell extracts were collected in lysis buffer (50 mM Tris-HCL, 150 mM NaCl, 2 mM EDTA, 1 % NP-40, 0,1% sodium dodecyl sulfate, 1 % sodium desoxycholate, 1 mM sodium orthovanadate, 1 mM phenylmethylsulfonyl fluoride, 50 mM sodium fluoride, 1x protease inhibitor cocktail (Cell Signaling, Beverly, MA, USA)). After sonication (EpiShear™ Probe sonicator, Active Motif, La Hulpe, Belgium) protein concentration was determined by the DC^TM^ Protein Assay (Bio-Rad Laboratories, Hercules, CA, USA). Afterwards, 1x Roti-Load was added. Tumor tissue samples (10-20 mg) were homogenized with Tissue Lyser (Qiagen, Hilden, Germany) in 300-600 µl 1x Roti^®^-Load buffer (Carl Roth GmbH, Karlsruhe, Germany). After sonication and centrifugation (10 min, 10.000 x g, 4 °C), the supernatant was removed. 20-35 µg of protein was denatured by heating (5 min, 95 °C) and separated by SDS-PAGE (6-15 % gels). Proteins were transferred to a nitrocellulose membrane (GE Healthcare, Little Chalfont, UK) via the Protean Mini Cell System (BioRad, München, Germany). The membrane was blocked with 5 % non-fat milk in TBS/0.1 % Tween 20 (MERCK, Darmstadt, Germany) (2 h, RT) and incubated with the corresponding primary antibody (1:500 to 1:10000, overnight, 4 °C). The following primary antibodies were used: HMOX1, N-Cadherin, pATM (S1981), pCHK2 (T68) and RAD51 from Abcam (Cambridge, UK), HIF1a, pKAP1 (S824) and pRPA32 (S4/S8) from Bethyl Laboratories Inc. (Montgomery, Texas, USA), BAX, BCL-2, cleaved Caspase-3, cleaved Caspase-7 (D198), Cyclin B1, Cyclin E1, GADD45A, GAPDH, P16 INK4A, P21 Waf1/Cip1, P27 Kip1, PARP, pCHK1 (S345), pMDM2 (S166), pP53 (S15), RhoA and Talin-1 from Cell Signaling Technology (Cambridge, UK). Cyclin D1 was purchased from Invitrogen (Darmstadt, Germany), γH2AX (S139) from Millipore (Billerica, MA, USA), ß-Actin, FASL, FASR, MKP-1 (DUSP1), P53 and Rac (pan) specific antibody from Santa Cruz Biotechnology (Santa Cruz, CA, USA), ERK2 from Thermo Fisher Scientific inc. (Waltham, MA, USA). After washing with TBS/0.1 % Tween 20, incubation with horseradish peroxidase-conjugated secondary antibodies (goat anti-mouse IgG and mouse anti-rabbit IgG (Rockland, Limerick, PA, USA)) was performed (1:2000, 2 h, RT). For chemiluminescence detection the ChemiDoc Imaging System (Bio-Rad Laboratories, Hercules, CA, USA) was used.

### Quantitative analyses of mRNA expression by RT-qPCR

Total RNA was purified from cells or 15-20 mg tumor tissue using the RNeasy Mini Kit (Qiagen, Hilden, Germany) according to the manufactureŕs protocol. The RNA yield was determined by measuring the OD260/280 ratio with NanoVue^TM^ Plus Spectrophotometer (GE Healthcare, Freiburg, Germany). RNA was isolated from 6-8 tumors isolated from 3-4 animals per experimental group and pooled for cDNA synthesis using the High-Capacity cDNA Reverse Transcription Kit (Applied Biosystem, Darmstadt, Germany). For each cDNA synthesis reaction, 2000 ng of total RNA was used. Quantitative real-time PCR analysis was accomplished with 20 ng of cDNA and 0.25 μM of the corresponding primers (Eurofins MWG Synthesis GmbH, Ebersberg, Germany) using CFX96 Touch Real-Time PCR Detection System (Bio-Rad Laboratories, Hercules, CA, USA). A semi-customized PCR array (Sigma-Aldrich, Steinheim, Germany) facilitating the analysis of the mRNA expression of genes related to cell-cycle regulation, DNA damage response (DDR), DNA repair, senescence, cell death and stress signaling as well as of inflammation, fibrosis and Rho GTPases-related factors or additional selected factors were used for quantitative RT-PCR analyses. Real-time qRT-PCR analyses were performed in technical duplicates or triplicates using a CFX96 cycler (Bio-Rad Laboratories, Hercules, CA, USA) and the SensiMix SYBR Kit (Bioline, London, UK). mRNA expression levels were normalized to the housekeeping genes ß-Actin and Gapdh. Quantitative RT-qPCR analysis was performed as follows: 1. 95 °C – 10 min; 2. 35-40 amplification cycles with 95 °C – 15 s, 55 °C – 15s and 72 °C – 17s; 3. 95 °C – 1 min, 55 °C – 1 min, 65 °C – 5 s. At the end of the run, we checked the melting curves to verify the correct product. PCR products with a cycle threshold >40 was excluded from the analysis. If not stated otherwise, the relative mRNA expression of non-irradiated controls (cells or tumors) was set to 1.0. Data were analysed using the CFX Manager Software (Bio-Rad Laboratories). Only changes in mRNA levels of ≤0.7- and ≥1.5-fold of the control were considered as biological relevant. Primers used for mRNA expression are from Eurofins (Ebersberg, Germany) and are listed in **Supplementary Table 1.**

### Animal experiments – syngeneic mouse model & fractionated in vivo irradiation

Female BALB/c mice (10-14 weeks old, body weight 20-30 g) were maintained in the central animal facility of the Heinrich-Heine-University Düsseldorf (ZETT). Animal experiments were conducted following the European Guidelines for the Care and Use of Laboratory Animals and were approved by the North Rhine-Westphalia State Agency for Nature, Environment and Consumer Protection (reference number: 81-02.04.2018.A305). 3-4 animals were used per experimental group. 1 × 10^5^ murine 4T1 cells were subcutaneously injected into the left and right flank of immunocompetent female mice. Body weight and tumor volume (tumor volume = length x width^2^/2) were recorded at least three times per week. When the tumor reached a volume of 0.3-0.5 cm^3^, local hFI of the tumors was performed (6 × 4 Gy; cumulative dose: 24 Gy; within two weeks) **(see Fig. 5A)**. Mice were anesthetized with ketamine (100 mg/kg bw) and xylazine (5 mg/kg bw) before irradiation with Gulmay RS 225 (15 mA, 200 kV) device. Tumors were locally irradiated in order to protect normal tissue from adverse effects using a self-constructed lead shield device ^63^. When the tumors reached a volume of ∼1 cm^3^, tumors and organs were isolated and either embedded in paraffin for immunohistochemical analyses or snap-frozen in liquid nitrogen and stored at −80 °C.

### Hematoxylin and eosin (H&E) staining

Formalin-fixed and paraffin-embedded primary tumor tissue was cut into sections of 4 µm thickness (RM 2164, Leica, Wetzlar, Germany). Paraffin removal and tissue rehydration were performed according to standard histological procedures. Afterwards, tissue sections were stained with hematoxylin/eosin (HE) (Sigma-Aldrich, Steinheim, Germany) and Sirius red (Waldeck, Münster, Germany) in order to evaluate morphological changes, inflammation, and fibrosis by light microscopy (Olympus BX43).

### Analysis of apoptosis by TUNEL assay

The frequency of apoptotic cells induced upon hFI *in vivo* was quantified by using the *in situ* Cell Death Detection Kit Fluorescein (TUNEL assay, Roche Diagnostics, Mannheim, Germany) according to the manufactureŕs instruction. Protein digestion was performed with 20 µg/ml Proteinase K (Qiagen, Hilden, Germany). Treatment with 150 U/ml DNase (Qiagen, Hilden, Germany) was performed as a positive control. Nuclei were counterstained with DAPI-containing Vectashield^®^ and analyzed with an Olympus BX43 microscope. 2-3 tumor sections obtained from 6-8 tumors that were isolated from 3-4 animals were analyzed in each experimental group and the percentage of TUNEL positive cells was calculated.

### Immunohistochemical analyses of tumor sections

Formalin-fixed and paraffin-embedded tumor tissue was cut into sections of 4 µm thickness. Xylol was used to remove paraffin and sections were rehydrated by ethanol/H_2_O according to the standard procedure. For antigen retrieval, rehydrated tumor tissue sections were incubated with Target Retrieval Solution (DAKO, Hamburg, Germany) in a steam boiler for 30 min, followed by blocking using Protein Block (DAKO, Hamburg, Germany) (1 h, RT). For immunohistochemical staining, tissue sections were incubated with primary antibody overnight (1:250 – 1:400; 4 °C; wet chamber). The following antibodies were used: rabbit anti-Ser139 phosphorylated histone 2AX (1:400, Abcam, Cambridge, MA, USA), rabbit anti-Ser10 phosphorylated histone 3 (1:250, Thermo Fisher Scientific). After washing (PBS/0.1% Tween 20 (3 × 5 min)) Alexa Fluor 488-coupled goat anti-rabbit secondary antibody (Invitrogen, Darmstadt, Germany) was used. Afterwards, sections were mounted in DAPI-containing Vectashield and evaluated with an Olympus BX43 microscope. Alternatively, avidin-biotin staining was employed for immunohistochemical analysis. To this end, sections were incubated with 3% H_2_O_2_ for 10 min to quench endogenous peroxidase activity. 0.001% avidin and 0.001% biotin were added for 15 min before primary antibodies were added (overnight, 4 °C). The following antibodies were used: rabbit anti-Ki-67 (1:400, Cell Signaling), rabbit anti-cleaved caspase 3 (1:300, Cell Signaling), rabbit anti-p21 (1:200, Cell Signaling), rabbit anti-CD3 (1:200, Abcam) to detect T-cells, rabbit anti-CD68 (1:100, Abcam) and mouse anti-MPO (1:500, R&D Systems), rabbit anti- *α*SMA (1:150, Abcam), rabbit anti-Vimentin (1:200, Cell Signaling) and mouse anti-CD144 (1:150, eBioscience). After washing, sections were incubated with biotinylated donkey anti-rabbit IgG (1:200, 45 min, RT, Santa Cruz, CA, USA). Afterwards, the sections were incubated with ABC reagent (Vectastain Elite ABC HRP Kit, Vector Laboratories, Burlingame, CA, USA) and stained with 3,3’-diaminobenzidine (DAB Peroxidase Substrate Kit (Vector Laboratories)) for 1-8 min. Sections were counterstained with hematoxylin, mounted in Entellan and evaluated using Olympus BX43 microscope. Tumor sections generated from 6-8 individual tumors isolated from 3-4 mice per experimental group were analyzed. Slides were microscopically evaluated using a 20x/40x objective.

### Statistical analyses

The unpaired, two-tailed Student’s t-test or the one-way ANOVA with Dunnett’s post-hoc test was used for statistical analysis. *p ≤ 0.05, **p ≤ 0.01, ***p ≤ 0.001 were considered as statistically significant differences between the groups. Quantitative data are shown as the mean ± SD or the mean ± SEM as clarified in the legends. Plotting of graphs and statistical analyses were performed by use of GraphPad Prism.

## Results

### 1. Comparative analyses of radiation-induced stress responses in parental (4T1^WT^) and radioselected (4T1^RS^) cells in vitro

To analyze whether repeated IR exposure of tumor cells leads to persisting alterations in the radiation response of the surviving progeny, murine mammary carcinoma cells (4T1) were exposed to a clinically oriented hFI protocol (4 Gy per irradiation) comprising a cumulative dose of 56 Gy applied over 6 weeks **(Fig. 1A)**. After a post-incubation period of 1.5 week, the surviving radioselected 4T1^RS^ cells were analyzed with respect to their response following a single IR exposure (4-8 Gy) as compared to parental (non-hFI-selected) 4T1 control (4T1^WT^).

#### 1.1. Analysis of proliferation cell cycle progression, and cell death

Both parental 4T1^WT^ and IR-selected 4T1^RS^ cells showed similar morphology **(Fig. 1A)** and revealed similar doubling time of approximately 24 h (**Supplementary Fig. S1**). For detailed comparative analysis of proliferation activity, EdU incorporation and mitotic index (pH3) were evaluated 24 - 72 h after a single IR with 4 Gy or 8 Gy. Both 4T1^WT^ and 4T1^RS^ cells showed a significant reduction in the percentage of EdU-positive cells following IR **(Fig. 1B and Supplementary Fig. S2A)**. However, the percentage of EdU-positive cells was significantly lower in irradiated 4T1^WT^ cells as compared to the radioselected 4T1^RS^ cells **(Fig. 1B and Supplementary Fig. S2A)**, indicating that 4T1^RS^ cells are more resistant to inhibition of S-phase activity following IR exposure than 4T1^WT^ cells. Analyzing the percentage of pH3-positive cells, 4T1^RS^ cells revealed a reduced mitotic index as compared to 4T1^WT^ under basal situation **(Fig. 1C).** Most important, a clear increase in the percentage of pH3-positive cells was observed in 4T1^WT^ cells after irradiation with 8 Gy, while the percentage of pH3 positive 4T1^RS^ cells remained unaffected **(Fig. 1C and Supplementary Fig. S2B)**. Overall, the data show that parental 4T1^WT^ cells are more sensitive to S-phase and mitosis-related toxicity evoked by single IR treatment than hFI selected 4T1^RS^ cells.

Next, we monitored the cell cycle distribution of 4T1^WT^ and 4T1^RS^ cells under basal situation and 24 - 72 h following irradiation with 4 Gy and 8 Gy by flow cytometry. 4T1^WT^ and 4T1^RS^ cells revealed a similar frequency of cells present in the SubG1- and G2/M-phase of the cell cycle under basal (non-irradiated) situation (**Fig. 2A** and **Supplementary Fig. S3A**), indicating that hFI does not provoke noticeable long-lasting effects on cell cycle distribution under basal situation. However, following a single IR treatment, distinct differences were observed between 4T1^WT^ and 4T1^RS^ cells. As analyzed at earlier time point (i.e., 24 h) after single IR exposure, 4T1^RS^ cells were characterized by a significantly higher percentage of cells present in G1-phase as compared to 4T1^WT^ cells **(Fig. 2A)**, while the percentage of 4T1^RS^ cells in G2/M phase was reduced **(Fig. 2A)**. The data are indicative of differences in the activation of G1/S-and G2/M-checkpoint mechanisms between parental 4T1^WT^ and radioselected 4T1^RS^ cells, supporting the hypothesis that hFI promotes persisting changes in cell cycle checkpoint-related mechanisms. At later time points (i.e., 48 h and 72 h), irradiated 4T1^WT^ revealed a significantly higher percentage of cells in the SubG1 fraction as compared to 4T1^RS^ cells **(Fig. 2B and Supplementary Fig. S3A),** indicating the hFI evokes acquired RR by protecting cells from IR-induced apoptosis. To confirm these data, the cleavage of PARP-1 and pro-caspases 3 and 7 were analyzed up to 72 h after irradiation with 8 Gy by western blot analyses. As shown in **Fig. 2C**, PARP-1 cleavage was detectable 72 h after IR treatment and was more pronounced in 4T1^WT^ than in 4T1^RS^ cells. Activation of pro-caspases also occurred at late time point after irradiation (i.e., 72h). Yet, there was no major difference detectable between parental cells and hFI selected variants **(Fig. 2C)**.

**Figure 2:**
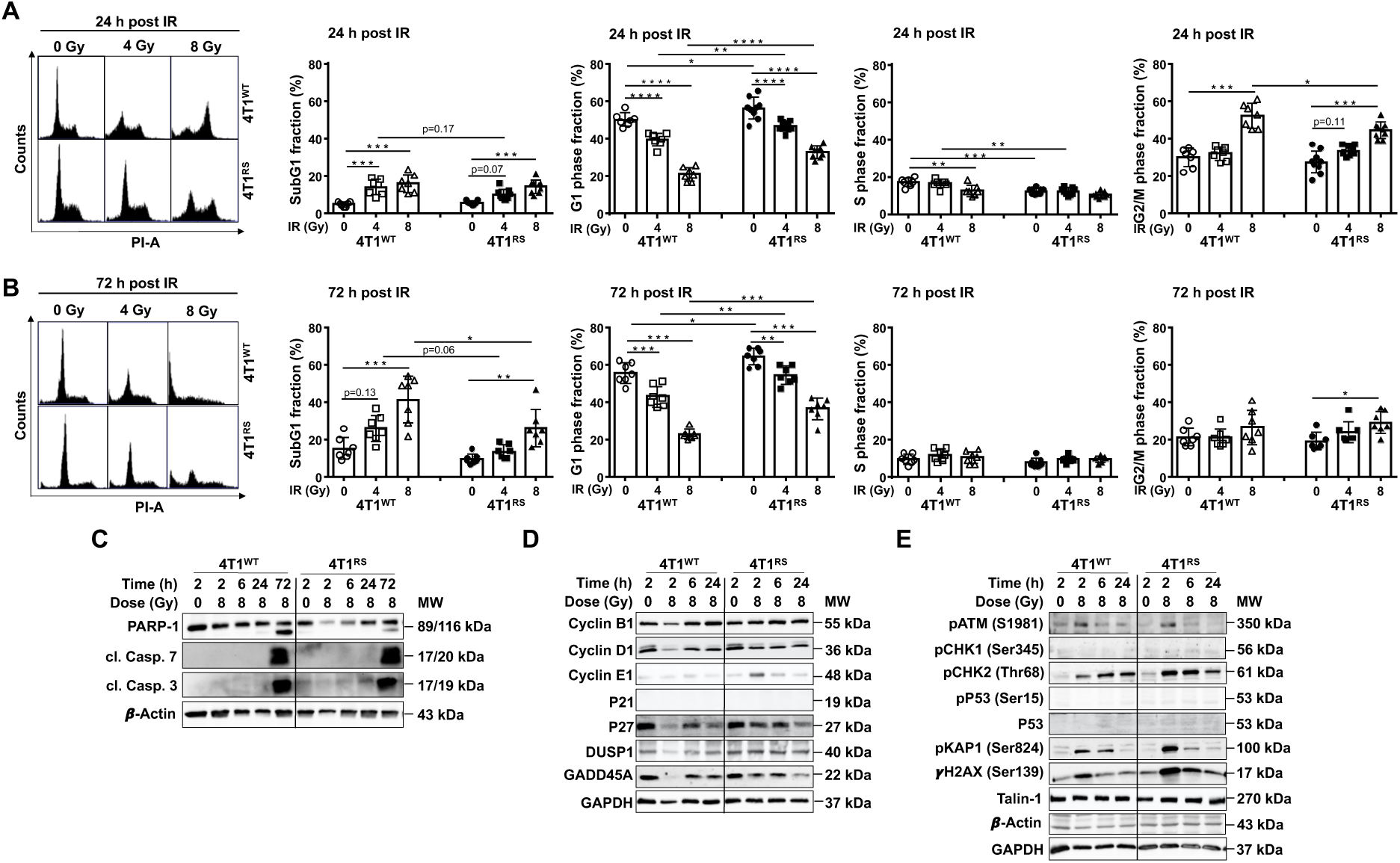
Cell cycle progression following ionizing radiation. **A-B:** Logarithmically growing 4T1^WT^ and 4T1^RS^ cells were irradiated with 4-8 Gy. After a post-incubation time of 24-72 h, cell cycle distribution was monitored by a flow cytometry-based method. The percentage of cells present in SubG1 fraction, G1-phase, S-phase, and G2/M-phase is shown. Quantitative data shown are the mean + SD from n=7-9 independent experiments each performed in biological duplicates. Statistical analysis was performed using one-way ANOVA with Dunnett’s post-hoc test. *, p ≤ 0.05; **, p ≤ 0.01; ***, p ≤ 0.001. Quantitative data obtained from the 48 h time point are shown in **Supplementary Fig. S2**. **C-E:** Logarithmically growing 4T1^WT^ and 4T1^RS^ cells were irradiated with 8 Gy. Protein extracts were collected 2 h, 6 h, and 24 h after treatment for western blot analysis. **(C)**: Shown are the protein levels of PARP-1 and cleaved Caspase-7 and −3. Expression of β-Actin was used as a loading control. **(D):** Shown are the protein levels of cyclin B1, cyclin D1, cyclin E1, P21, P27, DUSP1 and GADD45A. Expression of GAPDH was used as a loading control. **(E)** Shown are the protein levels of DDR-related factors: Ser1981 phosphorylated ATM (pATM), Ser345 phosphorylated checkpoint kinase-1 (pCHK1), Thr68 phosphorylated checkpoint kinase-2 (pCHK2), Ser15 phosphorylated protein 53 (pP53), Ser824 phosphorylated KRAB-associated protein-1 (pKAP1) and Ser139 phosphorylated histone 2AX (γH2AX). Expression of β-Actin, GAPDH and Talin-1 were used as loading controls.

In order to elucidate the molecular mechanisms underlying the differential activation of G1/S-and G2/M-checkpoint control mechanisms in 4T1^WT^ and 4T1^RS^ cells, protein expression of cell cycle regulatory proteins and DDR-related factors was analyzed by western blot analysis. The protein levels of G1-phase-related cyclin D1 and G2/M-phase-related cyclin B1 protein expression were transiently decreased in 4T1^WT^ cells 2 h after IR **(Fig. 2D).** 24 h after irradiation cyclin B1 level was slightly elevated in 4T1^WT^ as compared to 4T1^RS^ cells **(Fig. 2D)**. Interestingly, a transient increase in G1/S-phase-related cyclin E1 expression was observed 2 h after IR only in 4T1^RS^ cells **(Fig. 2D)**. The CDK inhibitory protein P21, which is a major target of p53 regulation, was not expressed in the p53 deficient 4T1 cells as expected ^64^. However, the CDK inhibitor P27, which also regulates cell cycle progression at pre-replicative G1-phase, was reduced with increasing time after IR exposure in both 4T1^WT^ and 4T1^RS^ cells (**Fig. 2D**). Expression of GADD45A, which interacts with various proteins involved in the regulation of gene expression and stress responses, including PCNA, p21, MEKK4 ^65–68^, was specifically diminished in 4T1^RS^ cells 24 h after IR exposure **(Fig. 2D)**. Regarding the activation of DDR-related factors we observed higher protein levels of phosphorylated pCHK2, pKAP1 and γH2AX at early time point (i.e. 2 h) after irradiation in hFI-selected 4T1^RS^ cells as compared to parental 4T1^WT^ cells **(Fig. 2E).** This data supports the hypothesis of long-lasting alterations of DDR-related stress responses going along with hFI. Having in mind the lack of functional P53 protein in 4T1 cells **(Fig. 2E)**, the persisting alterations in the responsiveness of 4T1^RS^ cells to IR-induced injury are obviously p53-independent.

#### 1.2. Comparative assessment of DNA damage formation, DNA repair and mRNA expression of susceptibility-related factors

To investigate the dose- and time-dependent formation of DNA damage induced by hFI, we performed immunocytochemical analyses of nuclear γH2AX foci and γH2AX/53BP1 co-localized foci as surrogate markers of DSBs. We found a significantly lower number of nuclear foci in 4T1^RS^ cells as compared with 4T1^WT^ cells as measured 2 h, 6 h and 24 h after treatment with 2-4 Gy. **(Fig. 3A and Supplementary Fig. S4)**. Furthermore, analyzing the IR-induced formation of micronuclei as additional marker of DNA damage (chromosome aberrations) ^69^, we observed a significantly lower number of micronucleated cells in 4T1^RS^ as compared to 4T1^WT^ cells (**Fig. 3B**). This data is indicative of accelerated repair of IR-induced DSBs and increased genetic stability in hFI-selected 4T1^RS^ cells.

**Figure 3:**
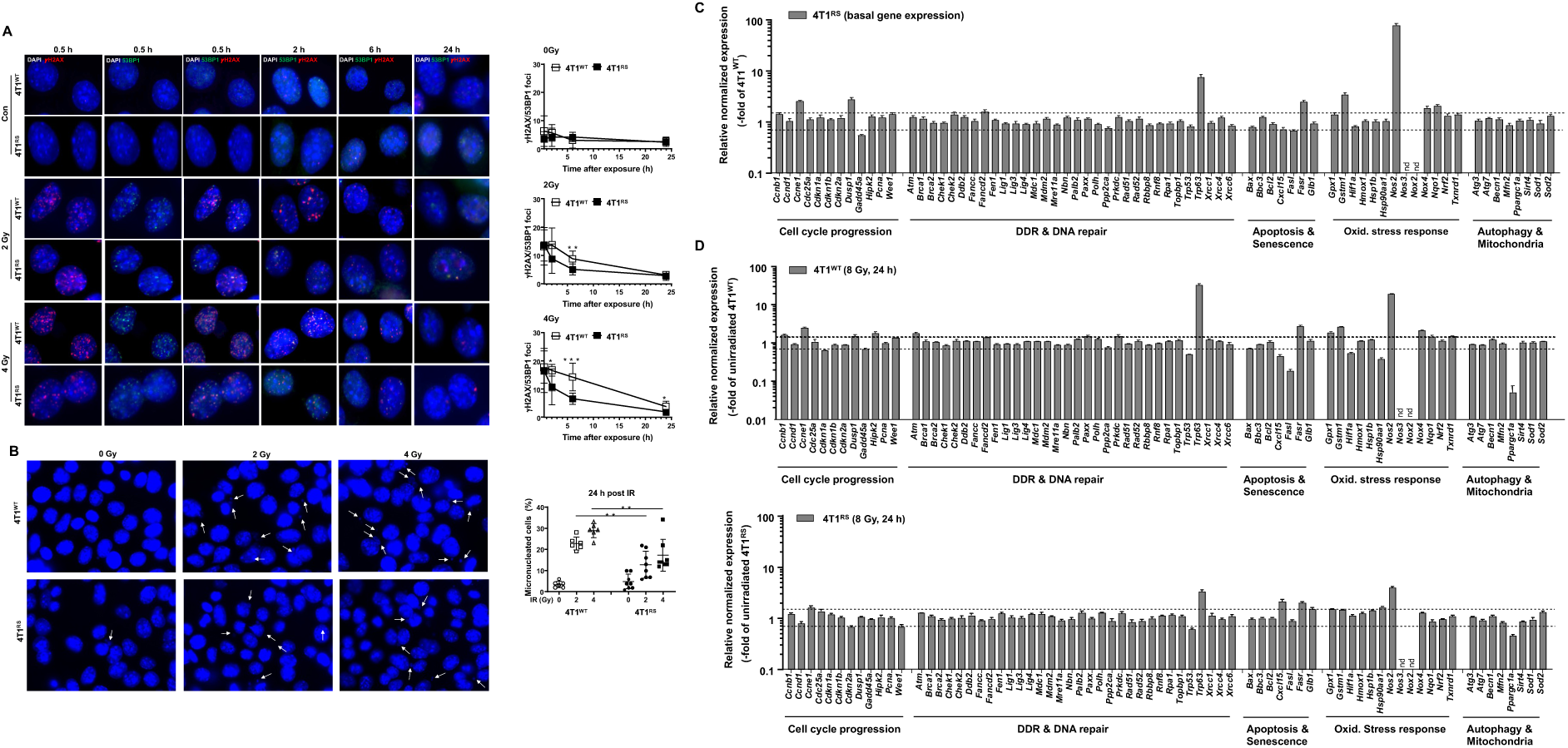
Formation and repair of IR-induced DNA damage. **A:** Logarithmically growing 4T1^WT^ and 4T1^RS^ cells were irradiated with 2-4 Gy. 30 min, 2 h, 6 h, and 24 h after treatment the formation of nuclear γH2AX foci and 53BP1 foci was analyzed as surrogate markers of IR-induced DNA double-strand breaks (DSBs) by immunocytochemistry. Representative images (left panel) and quantification (right panel) of γH2AX/53BP1 co-localized foci expressed as mean ± SD from n=4-5 independent experiments each performed in biological duplicates (N=6-10). ≥50 cells were analyzed per experimental condition. **B:** Logarithmically growing 4T1^WT^ and 4T1^RS^ cells were irradiated with 2-4 Gy. 24 h after irradiation the formation of micronucleated cells was analyzed. Representative images (left panel) and quantification (right panel) of micronucleated cells are shown. Quantitative data are shown as the mean ± SD from n=3-4 independent experiments each performed in biological duplicates (N=6-8). Statistical analysis was performed using unpaired, two-tailed Student’s t-test. *, p ≤ 0.05; **, p ≤ 0.01; ***, p ≤ 0.001 (irradiated cells as compared to the corresponding non-irradiated control). **C-D:** The mRNA expression of a subset of susceptibility-related genes was comparatively analyzed in 4T1^WT^ versus 4T1^RS^ cells under basal conditions **(C)** and **(D)** 24 h after IR treatment (8 Gy). Relative mRNA expression in parental non-irradiated 4T1^WT^ cells was set to 1.0. Changes in mRNA expression levels of ≥1.5 and ≤0.7 are considered as biologically relevant and marked with a dashed line. Data shown (mean ± SD) are technical duplicates obtained from pooled samples of biological triplicates.

To identify molecular mechanisms contributing to the phenotype of 4T1^RS^ cells, a comparative analysis of the mRNA expression of a selected subset of susceptibility-related factors, comprising factors regulating cell cycle progression, DNA damage response (DDR) and DNA repair, apoptosis, senescence, oxidative stress, autophagy and mitochondrial homeostasis, was performed. Under basal situation, we observed increased mRNA levels (≥1.5-fold) of *Ccne1, Dusp1, Trp63, Fasr, Gstm1, Nos2, Nox4, Nqo1* along with reduced mRNA levels (≤0.7-fold) of *Gadd45a and Fasl* in 4T1^RS^ as compared to 4T1^WT^ cells **(Fig. 3C)**. The most pronounced differences in mRNA expression levels between both models were observed for *Nos2* (70-fold) and *Trp63* (6-fold). Elevated basal mRNA expression of the Rho GTPases and Rho-regulatory factors *Rac3, Rhob, Arhgap1* and, most prominent, *Ncf2* (30-fold) were found in 4T1^RS^ cells, while the mRNA levels of *Ran1, Rhoc, Rhog, Pkn1* and *Tiam2* were reduced as compared to 4T1^WT^ cells (≤0.7-fold) **(Supplementary Fig. S5).** As measured 24 h after IR exposure (8 Gy) of 4T1^WT^ cells, substantial alterations in mRNA expression were detected for *Ccne1, Hipk2, p63, FasR, FasL, Gstm1, Nos2, Nox4* and *Ppargc1a (Pgc1α)* (**Fig. 3D, upper part**). Similarly, 4T1^RS^ cells revealed noticeable changes in the mRNA expression of *Ccne1, p63, FasR, Cxcl15, Nos2 and Ppargc1a (Pgc1α)* following exposure **(Fig. 3D, lower part)**. There were no major differences detectable between the IR-responsiveness of 4T1^WT^ and 4T1^RS^ cells. In summary, we conclude that hFI-selected 4T1^RS^ cells harbor specific and persisting changes in the basal mRNA expression profile of individual susceptibility-related genes as compared to parental 4T1^WT^ cells.

### 2. Effects of hypofractionated irradiation on in vivo growing 4T1 breast carcinoma cells

#### 2.1. Analyses of hFI on tumor growth, body and organ weight

Considering the potential influence of the tumor microenvironment on radiation responses of malignant cells, we next performed *in vivo* analyses utilizing a syngeneic immunocompetent xenograft mouse model. To this end, parental murine 4T1 mammary carcinoma cells were subcutaneously injected into the flank of BALB/c mice. 12 days after injection, the tumors formed were locally irradiated using a hFI protocol (6 × 4 Gy; 3 treatments per week) (IR-group (4T1^IR^)). A lead shield was used to protect normal tissue from adverse radiation effects **(Fig. 4A)** ^63^. As shown in **Fig. 4B**, hFI caused a transient delay in tumor growth up to day 28. Both non-irradiated tumors (4T1^CTR^) and tumors exposed to hFI (4T1^IR^) were isolated when they reached a volume of about 0.6-1.0 cm^3^ (4T1^CTR^: day 19; 4T1^IR^: day 31-33). No macroscopically detectable changes were observed between irradiated and non-irradiated tumors **(Fig. 4C, left panel)**. Yet, H&E staining of tumor sections revealed apoptotic and/or necrotic areas in the hFI-treated tumors as compared to non-irradiated tumors **(Fig. 4C, right panel)**. The body weight of non-irradiated and irradiated animals was similar **(Fig. 4D)**, showing that local hFI of tumor tissue (with simultaneous normal tissue shielding) does not cause noticeable systemic toxicity. Measuring organ weights, we observed a tendentially reduced liver weight and an elevated lung weight of the 4T1^IR^ mice as compared to the non-irradiated control **(Fig. 4E)**.

**Figure 4:**
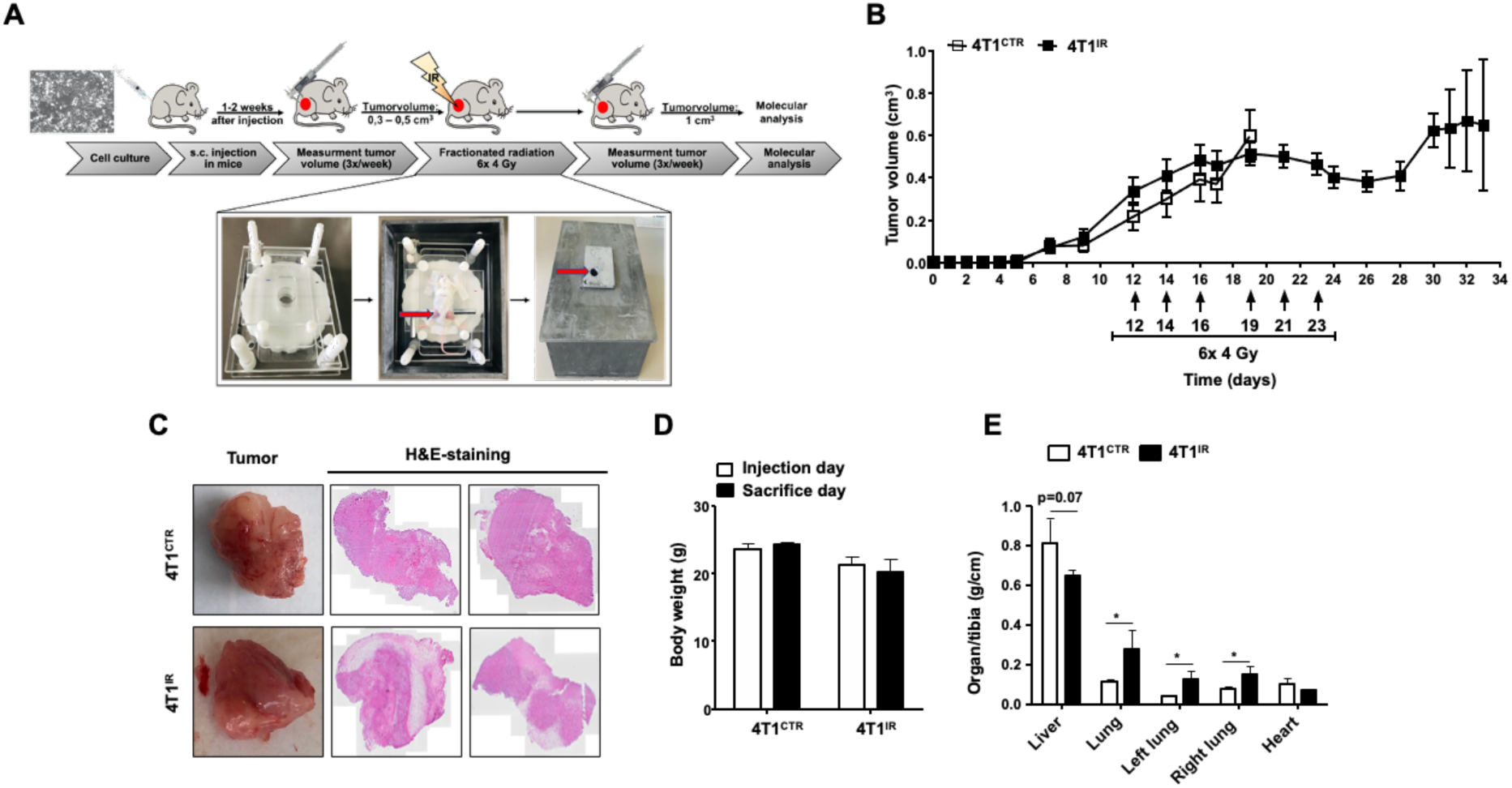
Hypofractionated irradiation of subcutaneously injected murine tumor 4T1 cells. **A, B:** Scheme of a clinically oriented hypofractionated irradiation protocol used. 10^5^ murine mammary tumor cells were injected into the left and right flanks of female Balb/c mice. The volume of the tumors was measured three times per week. When the tumors reached a size of 0.3-0.5 cm^3^, they were irradiated by placing the anesthetized mice in a self-made lead shield, which protects normal tissues from IR exposure. The animals were irradiated six times with 4 Gy (6 × 4 Gy) resulting in a cumulative dose of 24 Gy. Control animals were not irradiated. As soon as the tumors reached a volume of ∼1 cm^3^ the animals were sacrificed (non-irradiated 4T1^CTR^ tumors: on day 19; hFI-treated 4T1^IR^ tumors: day 28-33 after the s.c. tumor cell injection) for analyses tumor, liver, lung tissue. Data shown are the mean ± SEM of 4-8 tumors isolated from n=3-4 animals per experimental group. **C:** Representative images of macroscopical analysis of the tumor tissues (left) and corresponding H&E-staining of tumor tissue sections (20x magnification) (right). **D:** Analysis of body weight at the beginning and end of the experiment. Shown are the mean + SEM of n=3-4 animals per experimental group. **E:** Analysis of organ weight at the end of the experiment. Shown are the mean + SEM of n=3-4 animals per experimental group. Statistical analysis was performed using unpaired, two-tailed Student’s t-test. *, p ≤ 0.05.

#### 2.2. Effect of hypofractionated irradiation on tumor cell proliferation and apoptotic cell death in vivo

To investigate whether hFI causes persisting changes in the proliferation of *in vivo* growing tumors, mitotic index (pH3-positive cells) and the frequency of Ki-67 positive cells were analyzed. The data obtained show that the number of pH3-positive cells was not affected in the irradiated 4T1^IR^ tumors as compared to the 4T1^CTR^ tumors 10 days after the last hFI treatment **(Fig. 5A)**. By contrast, measuring the percentage of Ki-67 positive cells, we obtained a significantly reduced proliferative activity in irradiated 4T1^IR^ tumors as compared to the 4T1^CTR^ tumors **(Fig. 5B)**. This result demonstrates that hFI causes a sustained decrease in proliferation activity of 4T1^IR^ cells. Given that IR induces the formation of DNA damage, particularly DSBs, we evaluated the number of residual DSBs induced by hFI by analyzing the frequency of γH2AX positive cells. As shown in **Fig. 5C**, hFI enhanced the steady-state number of DSB harboring cells **(Fig. 5C)**. Apparently, DSBs induced by hFI are not yet fully repaired ten days after the end of the last exposure. In line with the fact that DSBs are potent inducers of cell death, we also observed a significantly increased number of TUNEL positive apoptotic cells in the irradiated 4T1^IR^ group **(Fig. 5D)** and a tendential increase in the percentage of cleaved Caspase-3-positive cells **(Fig. 5E).**

**Figure 5:**
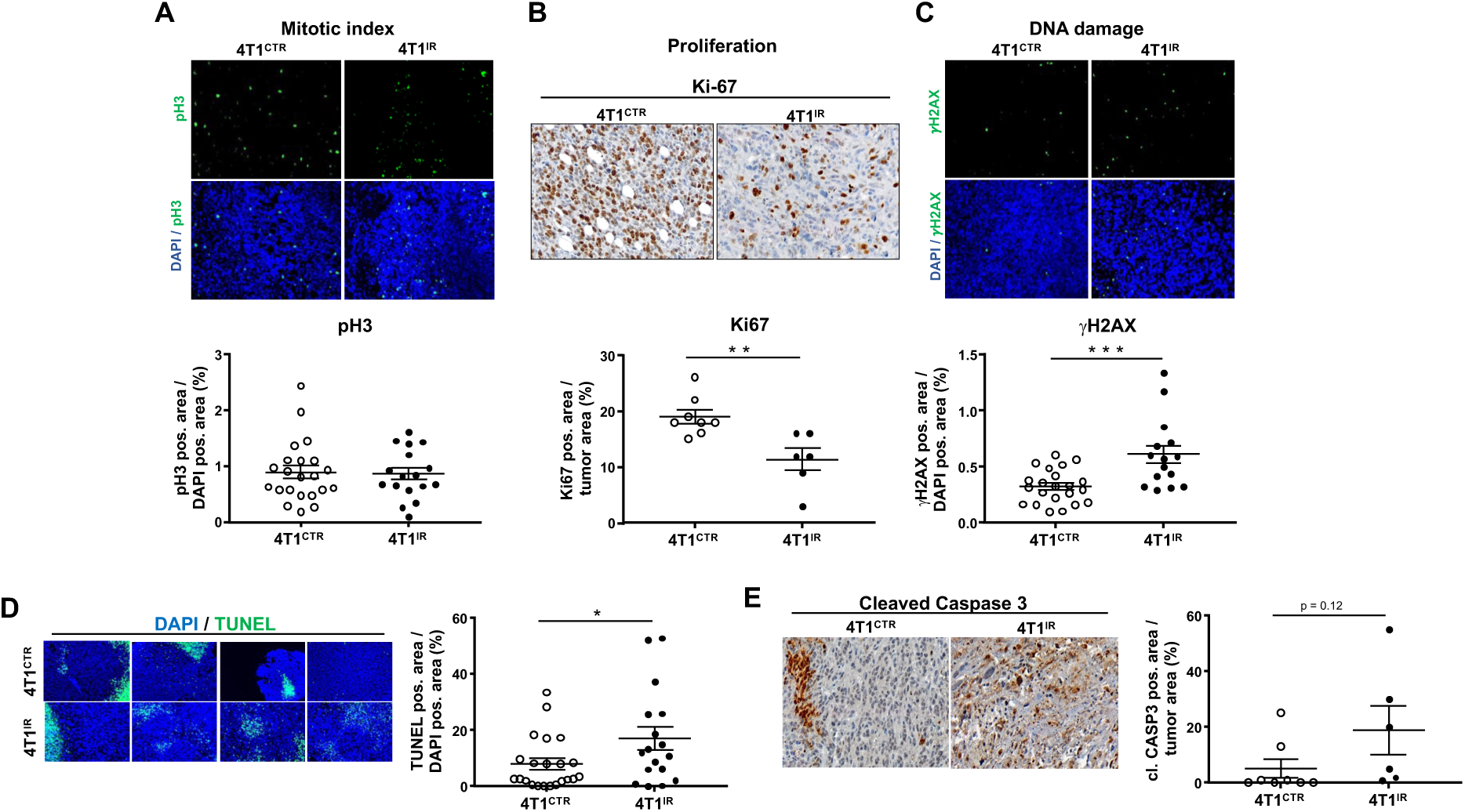
Analysis of proliferation, DNA damage and apoptosis in non-irradiated 4T1^CTR^ versus hFI-treated 4T1^IR^ tumors. **A-B:** Proliferation in non-irradiated 4T1^CTR^ and hFI-treated 4T1^IR^ tumors was analyzed by determination of the frequency of pH3-positive cells (mitotic index) (**A**), and Ki67-positive cells (**B**) in tumor sections as described in methods. Upper panel: Representative images of tumor tissue sections (20x magnification). Lower panel: Quantitative data are shown as mean ± SEM of 5-8 tumors isolated from n=3-4 animals per experimental group (biological duplicates in A). **, p ≤ 0.01. **C:** Residual DNA damage was investigated by the detection of nuclear γH2AX foci, which are indicative of DSBs. Upper panel: Representative images of tumor tissue sections (20x magnification). Lower panel: Quantitative data are shown as mean ± SEM of biological duplicates of 5-8 tumors isolated from n=3-4 animals per experimental group. ***, p ≤ 0.001 **D-E**: The frequency of apoptotic cells was determined by the TUNEL assay **(D)** and by immunohistochemical detection of cleaved Caspase 3-positive area **(E)** as described in methods. Nuclei are stained by DAPI (blue). Left panel: Representative images of tumor tissue sections (20x magnification). Right panel: Quantitative data are the mean ± SEM from 5-8 tumors (with one to three sections per tumor) isolated from n=3-4 animals per experimental group. *, p ≤ 0.05.

#### 2.3. Impact of hypofractionated irradiation (hFI) on the mRNA and protein expression of susceptibility-related genes in vivo

Next, we aimed to figure out whether hFI of *in vivo* growing 4T1 tumors causes prolonged (adaptive) alterations in the mRNA expression of genes involved in the regulation of cell cycle progression, DDR and DNA repair, cell death and senescence, oxidative stress or autophagy and mitochondrial-related functions. Regarding cell cycle regulatory factors, a reduced mRNA expression (≤ 0.7-fold) of *Cdkn1a and Gadd45a* was found in 4T1^IR^ as compared to 4T1^CTR^ tumors (**Fig. 6A**). Regarding DDR and DNA repair-related factors, we observed enhanced mRNA levels of the DSB repair-associated proteins *Rad51, Trp63* and *Xrcc6 (Ku70)* in 4T1^IR^ **(Fig. 6A)**. Moreover, 4T1^IR^ tumors revealed elevated mRNA levels (≥ 1.5-fold) of the oxidative stress-related factors *Gstm1, Nos2* and *Nqo1* as well as the mitochondrial proliferation-regulatory factor *Ppargc1a (PGC1α)* **(Fig. 6A)**. The mRNA levels of the cell death and senescence associated factor *Cxcl15* and oxidative stress-related factors *Hsp1b, Nos3, Nox2* and *Nrf2* was reduced in 4T1^IR^ as compared to 4T1^CTR^ tumors **(Fig. 6A)**. Overall, this data show that hFI results in persisting and complex alterations in the mRNA expression of multiple cell cycle-, DNA repair-, DDR-, cell death-, oxidative stress- and mitochondria-associated functions. Immunocytochemical analyses of cell cycle and senescence-related CDK-inhibitory protein p21 showed a tendential increase in the 4T1^IR^ tumors **(Fig. 6B)**. Analysis of the protein expression of representative susceptibility-related factors showed that protein expression of the G1-phase-related cyclin D1, G2/M-phase related cyclin B1 and GADD45A are reduced, while expression of G1/S-phase related cyclin E1, CDK inhibitor P27 and DUSP1 are elevated in the 4T1^IR^ tumors as compared to 4T1^CTR^ tumors (**Fig. 6C**). Furthermore, we found elevated protein levels of the DDR-related factors pCHK2, pP53 and γH2AX in 4T1^IR^ (**Fig. 6D**). Moreover, hFI triggered a sustained increase in the protein levels of cleaved PARP-1, cl. Casp7 and cl. Casp3 as well as FASR, and FASL (**Fig. 6E**). Additionally, an elevated protein level of HIF1α and a decreased protein amount of HMOX1 was found in 4T1^IR^ tumors (**Fig. 6F**). Taken together, we hypothesize that hFI of 4T1 mammary tumor cells growing in a syngeneic immunocompetent mouse model strengthens the development of G1/S checkpoint control mechanisms, likely promoting genomic stability and survival to hFI-induced damage.

**Figure 6:**
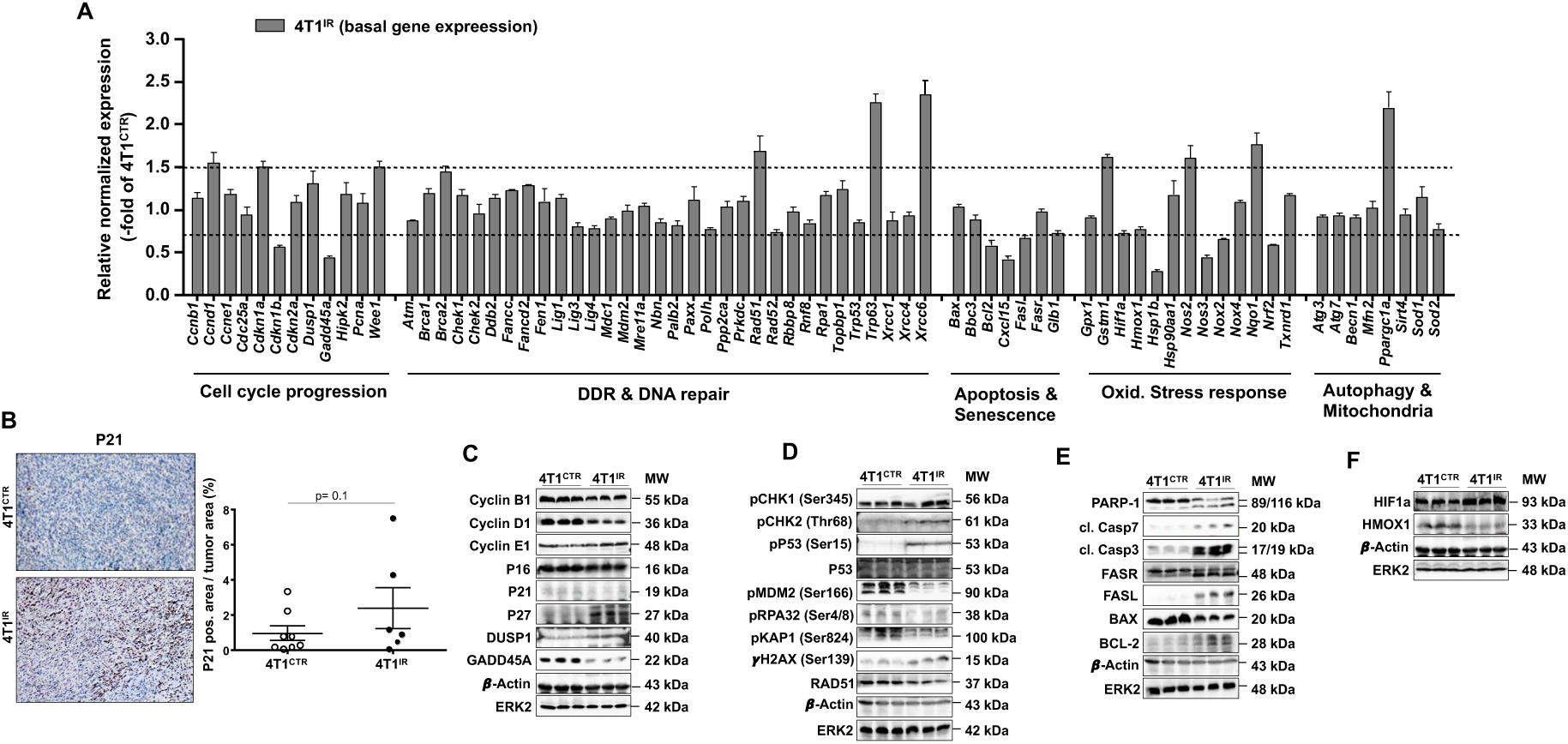
Comparative analysis of the expression of susceptibility-related factors. **A:** mRNA levels of susceptibility-related genes related to cell cycle progression, DNA damage response (DDR), DNA repair, apoptosis, senescence, oxidative stress response, autophagy, and mitochondria. Relative mRNA expression in 4T1^IR^ tumors was related to that of non-irradiated 4T1^CTR^ tumors, which was set to 1.0. Changes in mRNA expression levels of ≥1.5 and ≤0.7 are marked with dashed lines and are considered as biologically relevant. Data shown are the mean ± SEM from technical duplicates using pooled tissue samples from 6-8 tumors isolated from n=3-4 animals per experimental group. **B:** Immunohistochemical determination of the cell cycle and senescence-related marker P21. Left panel: Representative images. Right panel: Quantitative data are shown as mean ± SEM of tissue sections obtained from 5-8 tumors from 3-4 animals per experimental group. **C:** Analysis of the protein expression of cyclin B1, cyclin D1, cyclin E1, P16, P21, P27, DUSP1 and GADD45A in non-irradiated 4T1^CTR^ tumors and hFI-treated 4T1^IR^ tumors. Expression of β-Actin and ERK2 were used as protein loading controls. **D:** Analysis of the protein levels of representative DDR-related factors Ser345 phosphorylated checkpoint kinase-1 (pCHK1), Thr68 phosphorylated checkpoint kinase-2 (pCHK2), Ser15 phosphorylated protein 53 (pP53), protein 53 (P53), Ser166 phosphorylated MDM2 (pMDM2), Ser4/8 phosphorylated RPA32 (pRPA32) Ser824 phosphorylated KRAB-associated protein-1 (pKAP1), Ser139 phosphorylated histone 2AX (γH2AX) and RAD51 in non-irradiated control and irradiated tumors. Expression of β-Actin and ERK2 were used as protein loading controls. **E:** Analysis of the protein expression of apoptosis-related factors PARP-1, cl. Casp3, cl. Casp7, FASR, FASL, BAX and BCL-2. β-Actin and ERK2 were used as loading controls. **F:** Analysis of the protein expression of oxidative stress-related factors HIF1a and HMOX1 in non-irradiated and irradiated primary tumors. β-Actin and ERK2 were used as a loading controls.

#### 2.4. Impact of hypofractionated irradiation on inflammatory stress responses, immune cell infiltration, cytoskeleton-associated functions and fibrosis in vivo

A major reason for choosing a syngeneic mouse model for our analyses was that such a model allows the detection of hFI-induced stress responses related to the immune system. To investigate immune cell infiltration into tumor tissue, immunohistochemical (IHC) analyses were performed. The experiments revealed a significantly lower number of T-lymphocyte marker CD3-positive immune cells in 4T1^IR^ tumors as compared to the 4T1^CTR^ control **(Fig. 7A and 7B, upper part).** No differences in the infiltration of tumor tissue with macrophage marker CD68-positive cells (**Fig. 7A and 7B, middle part**), and a tendentially enhanced infiltration with neutrophil marker MPO-positive cells was observed in 4T1^IR^ tumors **(Fig. 7A and 7B, lower part)**. Furthermore, mRNA expression analyses of immune cell markers revealed largely enhanced mRNA levels of the lymphoid markers *CD19* (about 30-fold) *and a reduced mRNA expression of Ptprc (CD45) in* 4T1^IR^ **(Fig. 7C)**. *Regarding myeloid markers, increased mRNA levels of Il12a, Il10, Mpo, Ccl2 (MCP-1), Cxcl2,* and *Tnf* were found in 4T1^IR^ tumors, whereas the mRNA levels of *MhcII* and *F4/80* were reduced **(Fig. 7C)**. This data shows distinct prolonged immune system-related alterations in immune cell infiltration into malignant tissue occurring upon hFI *in vivo*.

**Figure 7:**
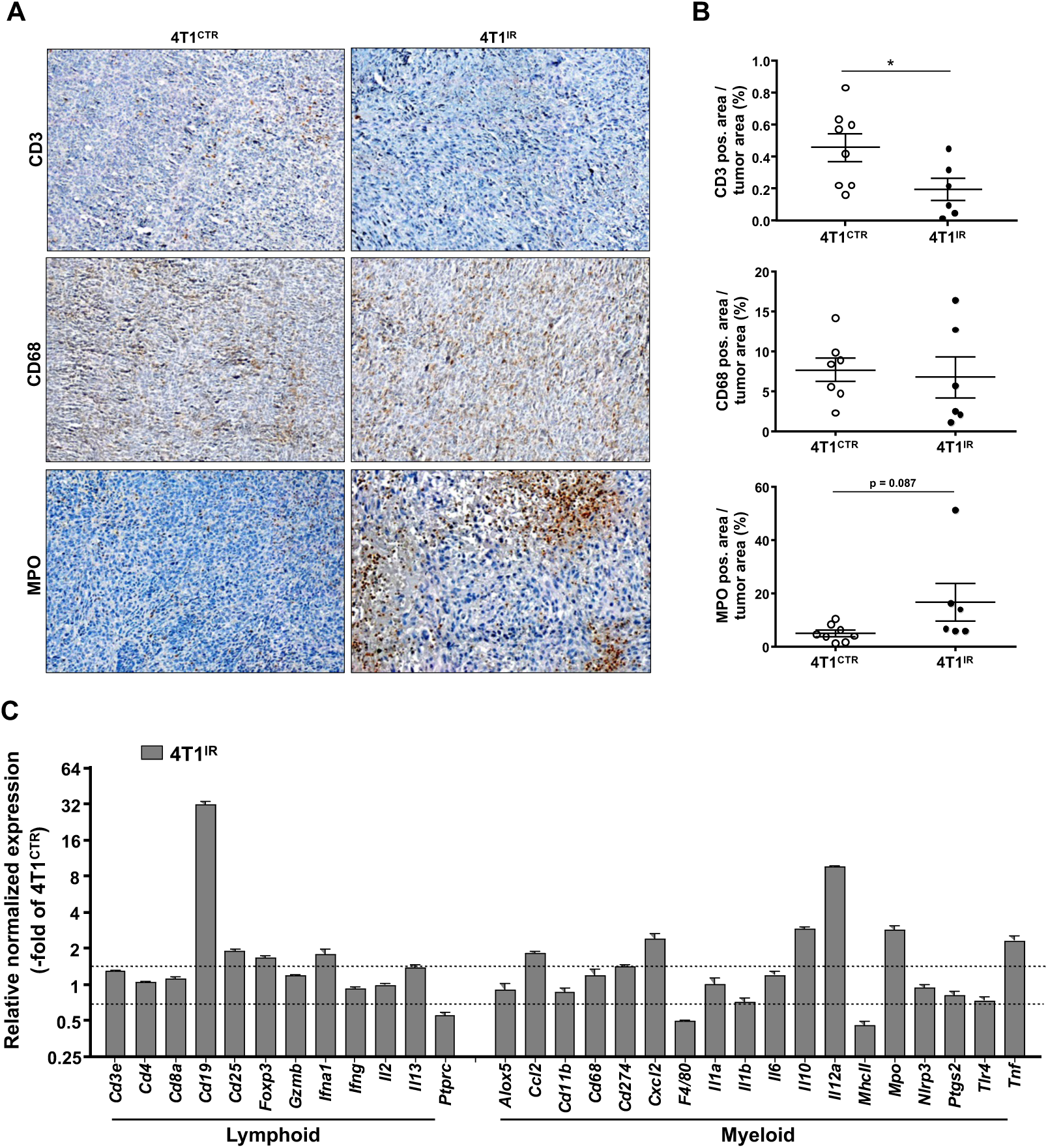
Influence of hFI on immune cell infiltration into 4T1^IR^ tumors. **A, B:** Immunohistochemical (IHC) staining of T-lymphocyte marker CD3, macrophage marker CD68 and marker of neutrophil MPO in tumor tissue sections of non-irradiated control (4T1^CTR^) and hFI irradiated tumors (4T1^IR^). Representative images (20x magnification) (**A**). Quantitative data are the mean ± SEM of 5-8 tumors isolated from 3-4 animals per experimental group (**B**). **C:** mRNA expression of lymphoid and myeloid marker genes was analyzed in 4T1^CTR^ and 4T1^IR^ tumors. Relative mRNA expression in irradiated 4T1^IR^ tumors was related to that of non-irradiated 4T1^CTR^ control, which was set to 1.0. Changes in mRNA expression levels of ≥1.5 and ≤0.7 as compared to the corresponding control are marked with dashed lines and are considered biologically relevant. Data shown are the mean ± SEM from technical duplicates using pooled tissue samples from 6-8 tumors isolated from n=3-4 animals per experimental group.

Additionally, the impact of hFI-induced effects on the expression of Ras-homologous (Rho) small GTPase, which are key regulators of the actin cytoskeleton and associated functions such as cell motility and cell adhesion ^70^ thereby affecting malignant transformation and metastasis ^71^was analyzed. We observed increased mRNA levels of the Rho GTPases *Rac2 and Rhot1 (Miro1)* as well as of Rho-regulatory GAPs and GEFs *Arhgap1 (Cdc42 GAP), Arhgap35 (p190RhoGAP), Racgap1, Rapgef3 (Epac1), Tiam2* (Rac GEF) and *Vav2 (Rac GEF)* **(Fig. 8A)**. By contrast the mRNA level of *Rhog* was reduced in 4T1^IR^ tumors. Overall, this data point to a so far unknown stimulatory effect of hFI on Rho GTPase-regulated functions, which play a key role in the regulation of tumor metastasis ^72–75^ and, most important, affects cancer radiotherapy ^76^. hFI-induced fibrotic tissue remodeling was analyzed by Sirius red staining of tumor sections. The hFI tumors showed increased thickness of mature collagen fibers in the Sirius Red staining of 4T1 tumors as compared to non-irradiated tumors (**Fig. 8B).** Moreover, immunohistochemical analysis revealed that the expression of the fibrosis-associated marker αSMA remains unchanged (**Fig. 8B**) and the expression of the cell adhesion protein CD144 (VE-Cadherin (CDH5)) increased in 4T1^IR^ tumors (**Fig. 8B**). Regarding the mRNA expression of tissue remodeling factors, we observed upregulated mRNA expression of collagenase *Col4a4*, *Ctgf* and matrix metalloproteases (*Mmp1, Mmp3,* and *Mmp9*), while the mRNA expression of *Col1a1, Col1a2, Col3a1 and Sftpc* were downregulated in the 4T1^IR^ as compared to 4T1^CTR^ tumors **(Fig. 8C)**. Analysis of cell-adhesion factors revealed elevated *Cd44* and *Icam1* mRNA levels, while the mRNA expression of *Bmp2, Cdh5 (VE-Cadherin, CD144), Cxcr4* and *Flk1* (*VEGFR-2*) was reduced in 4T1^IR^ tumors (**Fig. 8C**). Overall, this data demonstrates distinctive and sustained alterations in gene and protein expression of factors involved in the regulation of tissue remodeling and cell-cell adhesion, pointing to ongoing alterations in the tumor and/or the TME after the end of the hFI schedule *in vivo*.

**Figure 8:**
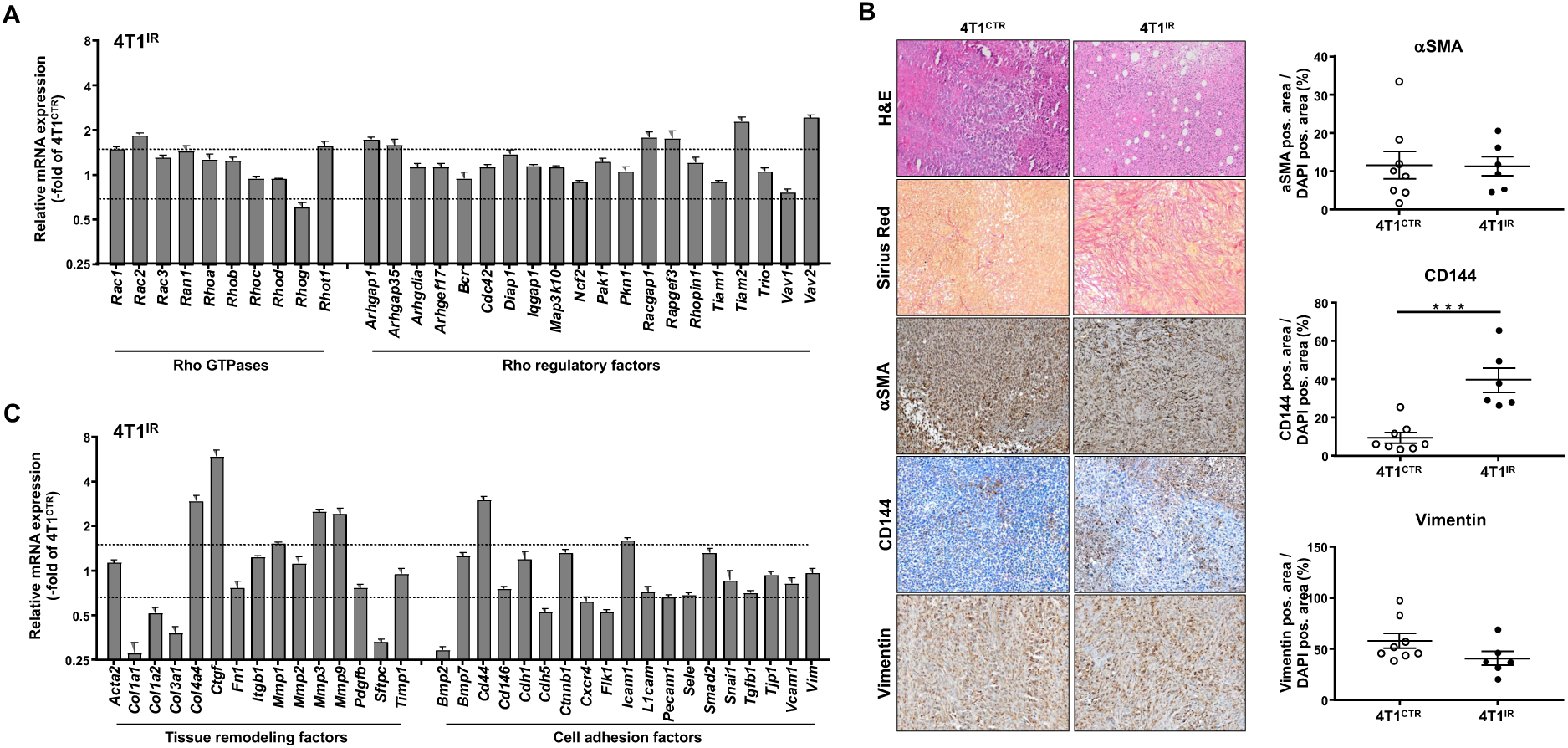
hFI-stimulated alterations in the expression of Rho GTPase-, tissue remodeling- and cell adhesion-related factors. **A:** mRNA expression of Rho GTPase and Rho-regulatory marker genes was analyzed in 4T1^CTR^ and 4T1^IR^ tumors. Relative changes in mRNA expression levels of ≥1.5 and ≤0.7 in 4T1^IR^ tumors as compared to the 4T1^CTR^ control, which was set to 1.0, are marked with dashed lines and are considered as biologically relevant. Data shown are the mean ± SEM from technical duplicates using pooled tissue samples from 6-8 tumors isolated from n=3-4 animals per experimental group. **B:** Immunohistochemical analysis of the protein expression of fibrosis and cell adhesion markers. Left panel: Representative images of tumor tissues sections stained by H&E, Picro-Sirius Red and results of immunohistochemical staining of the fibrosis marker αSMA, cell adhesion marker CD144 and EMT marker Vimentin (20x magnification). Immunohistochemical determination. Right panel: Quantitative data are shown as mean ± SEM of 5-8 tumors isolated from n=3-4 animals per experimental group. **C:** mRNA expression of tissue remodeling and cell adhesion-associated marker genes 4T1^CTR^ and 4T1^IR^. Relative changes in mRNA expression levels of ≥1.5 and ≤0.7 in 4T1^IR^ tumors as compared to the 4T1^CTR^ control, which was set to 1.0, are marked with dashed lines and are considered as biologically relevant. Data shown are the mean ± SEM from technical duplicates using pooled tissue samples from 6-8 tumors isolated from n=3-4 animals per experimental group.

#### 2.5. Comparative analysis of the mRNA expression of susceptibility related factors under in vitro versus in vivo situation

Finally, we aim to figure out whether the expression profile of susceptibility-related factors in non-irradiated 4T1 cells differs between *in vitro* growing 4T1 tumor cells (4T1^WT^) versus *in vivo* growing tumors (4T1^CTR^). Similarly, we aim to characterize changes in gene expression between *in vitro* radioselected cells (4T1^RS^) versus *in vivo* irradiated tumors (4T1^IR^). As compared to the *in vitro* growing non-irradiated 4T1^WT^ cells, the relative mRNA expression of susceptibility-related factors in the *in vivo* growing non-irradiated tumors (4T1^CTR^) was altered in a complex manner. As compared to their *in vitro* growing counterpart, *in vivo* growing 4T1^CTR^ tumors are characterized by an enhanced mRNA expression of individual genes involved in the regulation of cell cycle progression (i.e, *Cdkn1a, Dusp1a, Gadd45a, Hipk2*), DNA repair and DDR (*Trp63, Lig3, Lig4, Paxx*), apoptosis and senescence (*Bcl2, Glb1*), oxidative stress (*Nos2, Hmox1, Hsp1b, Hif1a*) as well as autophagy and mitochondrial functions (Atg7, *Sirt4*) **(Fig. 9A)**. By contrast, the mRNA expression of *Ccnb1, Chek1, Fancd2, Fen1, Lig1, Rad51, Topbp1* and *Nqo1* was found to be substantially downregulated in *in vivo* growing non-irradiated 4T1^CTR^ tumors **(Fig. 9A)**. Regarding the expression of Rho GTPases and Rho-regulatory factors, increase in the mRNA expression of *Ncf2, Vav1, Tiam2* and *Rac2* was observed, while mRNA levels of *Rac3, Ran1, Racgap1* and *Rhopin1* were reduced in 4T1^CTR^ tumors as compared to *in vitro* growing 4T1^WT^ cells **(Fig. 9B)**. The data show that the TME of *in vivo* growing 4T1 breast tumor cells selectively affects the basal mRNA expression of susceptibility-related and Rho-regulatory factors.

**Figure 9:**
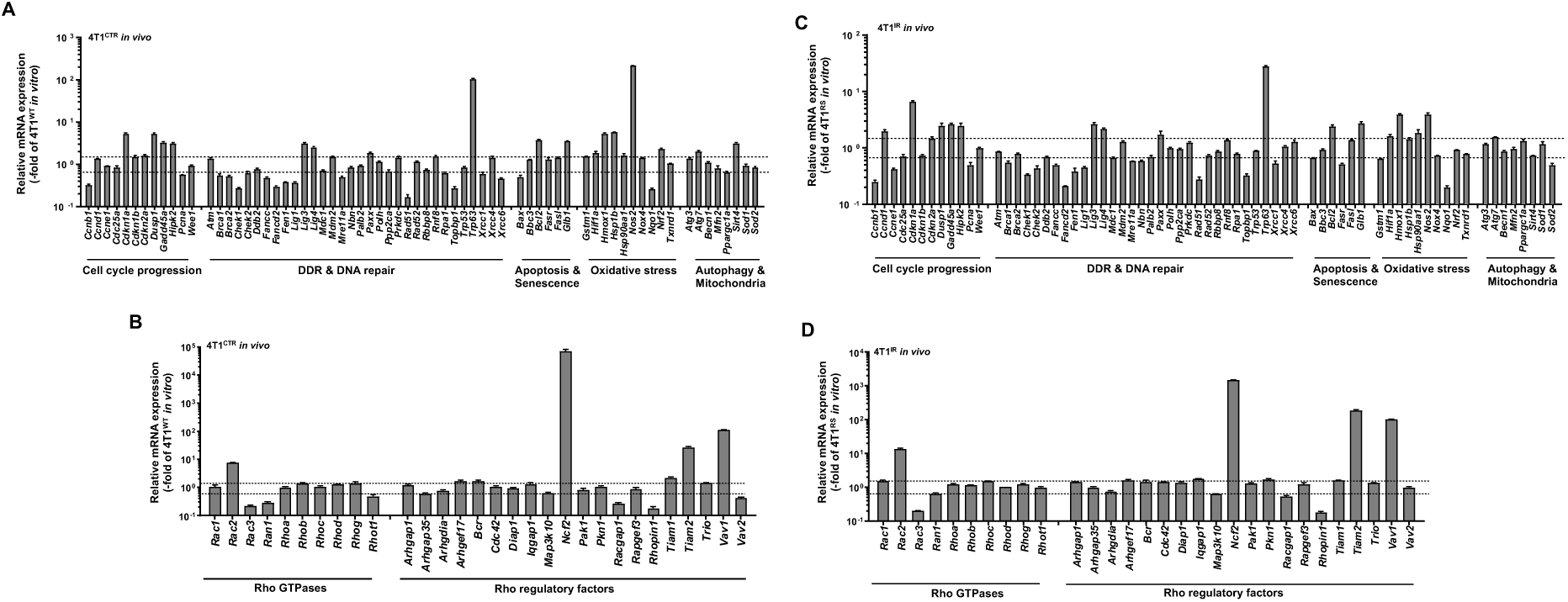
Comparative mRNA expression analyses of susceptibility-related factors and Rho-related factors under *in vitro* versus *in vivo* situation. **A, B**: mRNA expression of susceptibility factors **(A)** and Rho-related factors **(B)**. Shown is the relative mRNA expression in non-irradiated *in vivo* growing 4T1^CTR^ tumors as related to *in vitro* growing 4T1^WT^ cells, which was set to 1.0. Fold changes in mRNA expression levels under *in vivo* situation of ≥1.5 and ≤0.7 as compared to the corresponding *in vitro* control are marked with dashed lines and are considered biologically relevant. *In vitro* data shown are the mean ± SD from technical duplicates obtained from pooled samples of biological triplicates. *In vivo* are the mean ± SEM from technical duplicates using pooled tissue samples from 6-8 tumors isolated from n=3-4 animals per experimental group. **C, D:** mRNA expression of susceptibility factors **(C)** and Rho-related factors **(D)**. Shown is the relative mRNA expression in irradiated *in vivo* growing 4T1^IR^ tumors as related to *in vitro* growing 4T1^RS^ cells, which was set to 1.0. Fold changes in mRNA expression levels under *in vivo* situation of ≥1.5 and ≤0.7 as compared to the corresponding *in vitro* control are marked with dashed lines and are considered biologically relevant. *In vitro* and *in vivo* gene expression data were obtained as described above.

Similar, comparative analyses of hFI-inducible responses *in vitro* versus *in vivo* revealed elevated hFI-stimulated expression levels of genes involved cell cycle progression (i.e, *Cdkn1a, Dusp1, Gadd45a, Hipk2*), DNA repair and DDR (*Trp63, Lig3, Lig4, Paxx*), apoptosis and senescence (*Bcl2, Glb1*), oxidative stress (*Nos2, Hmox1, Hsp1b, Hif1a*) and autophagy and mitochondrial functions (*Atg7*) in 4T1^IR^ tumors as compared to 4T1^RS^ cells **(Fig. 9C)**. By contrast, the mRNA expression of *Ccnb1, Ccne1, Chek1, Fancd2, Fen1, Lig1, Rad51, Topbp1* and *Nqo1* was found to be substantially downregulated in *in vivo* growing irradiated 4T1^IR^ tumors **(Fig. 9C)**. Regarding the expression of Rho GTPases and Rho-regulatory factors in response to hFI, increased mRNA expression of *Ncf2, Vav1, Tiam2* and *Rac2* was observed while mRNA levels of *Rac3, Ran1, Racgap1* and *Rhopin1* were reduced in 4T1^IR^ tumors as compared to *in vitro* growing irradiated 4T1^RS^ cells **(Fig. 9D)**. Overall, this data, which are graphically summarized in **Supplementary Fig. S6**, show that the TME of *in vivo* growing tumor cells selectively impacts the mRNA expression of susceptibility-related and Rho-regulatory factors under situation of hFI.

## Discussion

The aim of this study was to identify molecular mechanisms that contribute to the development of a radioresistant phenotype in malignant cells following *in vitro* and *in vivo* treatment of murine mammary breast cancer cells using a clinically oriented hFI protocol. Under *in vitro* conditions, radioselected 4T1^RS^ cells were isolated after hFI (cumulative dose of 56 Gy) and compared with non-irradiated parental 4T1^WT^ cells. The radio-selected 4T1^RS^ cells exhibit persistent changes in IR-inducible cell cycle-associated mechanism related to S-phase activity, as analyzed by measuring EdU incorporation, G1/S checkpoint activation as analyzed by flow cytometry, a transiently altered expression of cyclins as well as a pronounced DNA damage response (DDR). These complex persisting adaptations observed in the radioselected 4T1^RS^ cells are accompanied by reduced PARP cleavage and a lower percentage of SubG1, indicating that their RR is due to an attenuated radiation-induced apoptosis as compared to parental 4T1^WT^ cells. A key factor of the DDR, which is well known to regulate both cell cycle progression, DDR and apoptosis, is p53. However, the 4T1 cell model used is p53 defective ^64^. In line with this, expression of p53 protein and p53-regulated factor p21 was not detectable, neither in the wild-type nor the hFI-selected cells. Accordingly, we concluded that the radioresistant phenotype of 4T1^RS^ is independent of p53-regulated functions. We hypothesize that hFI ultimately triggers the activation of adaptive mechanisms in 4T1^RS^ cells that compensate for their p53 deficiency, especially with respect to cell cycle checkpoint control mechanisms, including changes in the radiation-induced G1/S block that contributes to radioresistance ^77^. In line with this hypothesis, irradiated wild-type 4T1^WT^ cells showed a transiently reduced cyclin B1 and D1 protein expression as well as a transient reduction in the expression of p27 and Gadd45a protein. Noteworthy, Gadd45a interacts with cyclin B1 and D1 and is involved in the regulation of cell cycle checkpoints in the S phase and G2/M arrest ^65, 78^. In addition, we speculate that the increase in pH3-positive 4T1^WT^ cells observed after single irradiation is mediated by Aurora B kinase ^79, 80^, which is relevant for chromosome segregation during mitosis ^81, 82^. Because this effect is no longer observed in 4T1^RS^ cells, we hypothesize that hFI promotes the dephosphorylation of pH3, for instance by activation of phosphatase PP1 ^80^ and DUSP1 ^83^, which is known to mediate radioprotective effects ^84^. Consistent with this hypothesis, we observed an increased DUSP1 protein expression in 4T1^RS^ cells compared to 4T1^WT^ cells. Summarizing, we suggest that the hypofractionated *in vitro* radiation regimen triggers complex and prolonged adaptive changes of cell cycle regulation-related mechanisms, which ultimately increase the proliferative activity of the radioselected 4T1^RS^ cells, thus leading to a radioresistant phenotype.

Given that DNA double-strand breaks (DSBs) are the most cytotoxic DNA lesions induced by IR, it is feasible that DSB repair is also changed in 4T1^RS^ cells. Indeed, we observed a faster repair of IR-induced DSBs and a lower number of residual DSBs in the radioselected 4T1^RS^ cells. Accordingly, we assume that RR acquired through hFI is at least partially due to a more efficient repair of radiation-induced DSB. Since changes in homologous recombination repair (HR) that is taking place in S- and G2-phase, are possible mechanisms of RR ^85^, we speculate that improved pre-replicative repair of IR-induced DSB formation in 4T1^RS^ cells together with a more pronounced activation of DDR-related pro-survival mechanisms contributes to their RR phenotype. For further clarification, gene expression of selected susceptibility-related factors was investigated under basal situation and 24 h after single irradiation of wild-type and radioresistant cells. The results of these studies show complex differences in the both basal and IR-induced mRNA expression of genes involved in cell cycle regulation, DNA repair and DDR regulation, apoptosis and senescence, as well as autophagy and oxidative stress between wild-type and radioselected cells. Since 4T1 cells are p53-deficient, the observed alterations in the mRNA expression, including that of known p53 target genes such as p21, Bax, or Fas receptor, are obviously due to p53-independent mechanisms. In summary, we show that hFI of murine mammary carcinoma cell *in vitro* leads to RR that is likely attributable to multiple p53-independent mechanisms associated with the regulation of cell cycle progression, cell death and DNA repair. Based on literature, it is tempting to speculate that the adaptive mechanisms involved comprise other members of the p53 family, in particular p63 ^86^.

In a next step, a syngeneic mouse xenograft model was employed to investigate the effects of hFI *in vivo*. These analyses aimed to identify similarities and/or differences between hIF-inducible responses of p53-deficient murine mammary carcinoma cells *in vitro* versus *in vivo*. Based on theoretical considerations, we hypothesize that HFI-triggered stress responses of tumor cells *in vitro* differ from those *in vivo* because of the tumor microenvironment (TME), which is known to impact the growth and stress responses of tumor cells *in vivo* ^53, 87^. As expected, the *in vivo* irradiated tumors (4T1^IR^) showed a significantly reduced expression of the proliferation marker Ki-67 as compared to the corresponding non-irradiated controls (4T1^CTR^), but not of the mitotic index marker pH3, indicating that a large fraction of 4T1^IR^ cells are in G0/G1 phase and quiescent ^88^. In addition, the irradiated tumor cells revealed a higher number of residual DSBs than non-irradiated 4T1^CTR^ as well as an elevated frequency of TUNEL- and cleaved caspase 3-positive cells, pointing to ongoing apoptotic processes. Since the mRNA expression of the p53 target genes Bax and FAS receptor was not enhanced, we assume that hFI-simulates caspase-regulated apoptosis in 4T1^IR^ cells, which is independent of the p53-BAX/FASR axis. Rather, elevated protein expression of FASL, which is regulated in an AP1-dependent manner ^89^ might be involved. Noteworthy, a reduced protein expression of cyclin B1, cyclin D1, and GADD45a was observed in the irradiated 4T1^IR^ tumors *in vivo*, which is in accordance with what we observed in 4T1^RS^ cells *in vitro*. Apparently, cell cycle-related hFI-induced stress responses are conserved *in vitro* and under *in vivo* conditions. So, it is feasible that the reduced RNA and protein expression of *Gadd45a* evoked by hFI contributes to an increased induction of cell death as suggested by others ^90, 91^. Noteworthy in this context, TAp63, that is regulated in a NF-κB dependent manner and involved in cell cycle regulation, differentiation and survival upon NF-κB activation by multiple stimuli ^92^, has no influence on *Gadd45a* expression ^90^. We speculate that the downregulation of GADD45a following hFI disrupts genomic stability and G2/M checkpoint control mechanisms and suppresses apoptosis ^93, 94^. The observed alterations in the protein expression of DUSP1 phosphatase may also contribute to the development of RR in tumors ^84^. Mechanistically, the upregulation of DUSP1 may protect from hFI-induced apoptosis by inhibiting radiation-stimulated pro-apoptotic JNK1 activity ^95^. Furthermore, the protein expression of pMDM2, which regulates the turnover of p53 ^96^, was lower in 4T1^IR^ as compared to 4T1^CTR^ tumors. This indicates that the stability of p53 family members, such as p63 and p73, which are important in tumor development ^97, 98^, is affected upon hFI *in vivo*. Noteworthy, the mRNA expression of *p63* was elevated in 4T1^IR^, supporting the hypothesis of hFI-induced ongoing alterations of p63-regulated mechanisms in 4T1^IR^ tumors. Apart from p63, the mRNA expression of *Xrcc6/Ku70* and *Rad51* was upregulated in 4T1^IR^ tumors, indicating that hFI fosters DSB repair by NHEJ and HR *in vivo*. Another interesting observation is that the mRNA expression of *Ppargc1a*, which is a key regulator of mitochondrial proliferation ^99^ was upregulated in 4T1^IR^ tumors. This finding supports the view that, apart from genomic DNA damage, hFI causes mitochondrial DNA damage that disturbs mitochondrial homeostasis, thus promoting alterations in tumor cell metabolism. In line with this hypothesis, release of mitochondrial DNA and related activation of the cGAS/STING pathway has been reported upon IR treatment ^100^.

It is well known that the expression of immune system-related factors changes in locally growing tumors after radiotherapy, that is indicative of an inflammatory TME ^101–103^. In line with this, the results of our immunohistochemical studies revealed a tendential increase in the MPO-positive tumor area, while the CD3-positive area was reduced. Moreover, we observed a largely increased mRNA expression of the lymphoid marker *Cd19* and several myeloid markers, including *Il12a, Il10, Mpo* and *Cxcl2* in 4T1^IR^ tumors as compared to non-irradiated 4T1^CTR^ control. This data point to an enhanced infiltration of 4T1^IR^ tumors with B-lymphocytes, monocytes and neutrophils. By contrast, 4T1^IR^ tumors reduced mRNA levels of *Ptprc/CD45, F4/80* and *MhcII* are indicative of alterations in T- and B-cell antigen receptor signaling, macrophage infiltration and adaptive immune responses, respectively. Thus, our data highlight complex alteration in tumor immunology occurring in response to hFI *in vivo*.

The expression of many inflammatory cytokines is NF-κB dependent and regulated in a Rho (Ras-homologous)-dependent manner ^104, 105^. Moreover, Rho GTPases play a key role in the regulation of the actin cytoskeleton, tumor development, invasion and metastasis as well as proliferation and apoptosis ^106, 107^. Accordingly, the question arose whether the expression of Rho-associated factors change upon hFI. Analyzing the mRNA expression of Rho GTPases and Rho-regulatory factors, we found increased mRNA expression of Rho GEFs (*Rapgef3, Tiam2, and Vav2*) and Rho GAPs (*Arhgap1, Arhgap35, and Racgap1*) in 4T1^IR^ tumors. Noteworthy, the Rho GTPase Rac1 mediates RR in mammary carcinoma cells ^108, 109^ and, furthermore, high levels of Rac1 promote the survival of breast cancer cells following FI *in vitro* ^110^. Having in mind that high levels of Rac1 were associated with poor prognosis ^111^, Rac1 may be a useful prognostic and therapeutic target in radioresistant breast cancer cells. Overall, our findings provide evidence that hFI leads to a complex change in Rho/Rac-regulated signaling mechanisms *in vitro* and *in vivo* that are involved in the regulation of RR and metastasis-related processes. In this context, it should also be mentioned that epithelial-mesenchymal transition is regulated in a Rac-dependent manner ^112–114^. In line with this report, we found substantial changes in the expression of the tissue remodeling and fibrosis-associated factors *Col4a4, Ctgf, Mmp1, Mmp3, Mmp9* as well as the metastasis-related factor *Cd44* in 4T1^IR^ tumors, while the expression of *Col1a1, Col3a1, Sftpc, Bmp2* was reduced. Increased expression of the metastasis-associated factor CD144 (VE-Cadherin) was also found in immunohistochemical analyses.

Another aim of our study was to figure out whether the expression profile of susceptibility-related factors differs between *in vitro* growing 4T1 cells (4T1^WT^) versus *in vivo* growing tumors (4T1^CTR^) under basal situation or following hFI *in vitro* versus *in vivo* (i.e. 4T1^RS^ cells versus 4T1^IR^ tumors). The results of these extensive studies revealed that the *in vivo* growing 4T1^CTR^ cells are characterized by complex, either enhanced or reduced mRNA expression of individual genes involved in the regulation of cell cycle progression (e.g., *Cdkn1a, Dusp1, Gadd45a, Hipk2, Ccnb1*), DNA repair and DDR (*Trp63, Lig3, Chek1, Fancd2, Fen1, Lig1, Rad51, Topbp1*), apoptosis and senescence (*Bcl2, Glb1*), oxidative stress (*Nos2, Hmox1, Hsp1b, Nqo1*) and autophagy and mitochondrial functions (Sirt4). The same holds true with respect to the expression of Rho GTPases and Rho-regulatory factors (e.g., *Ncf2, Vav1, Tiam2, Rac2, Rac3, Ran1, Racgap1 and Rhopin1*). Similarly, some of the long-lasting hFI-induced transcriptional stress responses were also overlapping between *in vitro* versus *in vivo* growing cells, including genes involved the regulation of cell cycle progression, DNA repair and DDR, apoptosis and senescence, oxidative stress and autophagy, mitochondrial functions and Rho GTPases **(Supplementary Fig. S6)**. The data demonstrate that the TME of *in vivo* growing tumor cells selectively affects the mRNA expression of susceptibility factors and Rho-related factors both under non-irradiated basal situation and following hFI.

## Conclusion

Long-term hFI evokes adaptive changes in cell cycle checkpoint-related mechanisms and DSB repair that are facilitating DNA replication and the activation of anti-apoptotic pathways, eventually promoting RR. P53-deficiency of 4T1 breast cancer cells is likely compensated by other P53-family members to maintain RR. Moreover, hFI triggers the infiltration of myeloid and lymphoid immune cells into the tumor, likely fostering tumor progression. Based on the data we hypothesize that acquired RR is supported by mechanisms related to Rho GTPases, especially Rac1-related factors, GADD45A, DUSP1, CD44 and NQO1. Moreover, we hypothesize that hFI triggers ROS production, which in turn stimulates antioxidative stress responses further contributing to RR. Apart from tumor cell-specific responses to hFI, radiation-induced stress responses of the TME also contribute to RR under *in vivo* situation. Hence, future improvement of the anticancer efficacy of radiotherapy requires to consider both branches of radiation-induced responses and a more detailed dissection of persisting stress responses evoked by prolonged FI.

## Acknowledgement

The work was supported by the Wilhelm Sander foundation (AZ: 2018.018.1). We thank Lena Abbey and Claudia Gavranic for excellent technical support as well as Julia Mann and Marlena Sekeres for supporting animal experimentation.

## Abbreviations

ATM: ataxia telangiectasia mutated
CHK1/2: checkpoint kinase 1/2
DDR: DNA damage response
DSB: DNA double-strand breaks
DUSP: dual specificity phosphatase
hFI: hypofractionated irradiation
HIF: hypoxia-inducible factor
HMOX: heme oxygenase
γH2AX: Ser139 phosphorylated histone H2AX
GADD: growth arrest and DNA damage
IR: ionizing radiation
KAP1: KRAB associated protein 1
MDM2: murine double minute 2
MPO: myeloperoxidase
PARP: poly-ADP ribose polymerase
p53: tumor suppressor p53
p21: cyclin-dependent kinase inhibitor 1
pH3: phospho-histone H3
PCR: polymerase chain reaction
Rho: Ras-homologous
Rac: Ras-related C3 botulinum toxin substrate
RPA: replication protein A
RR: radioresistance
RT: radiotherapy
SMA: smooth muscle actin
4T1: murine mammary carcinoma cells.

