## Supplementary figures for "Comparative analyses of prolonged stress responses of breast carcinoma cells after hypofractionated irradiation *in vitro* and *in vivo*": Sivakumar et al_Supplementary Figures.docx

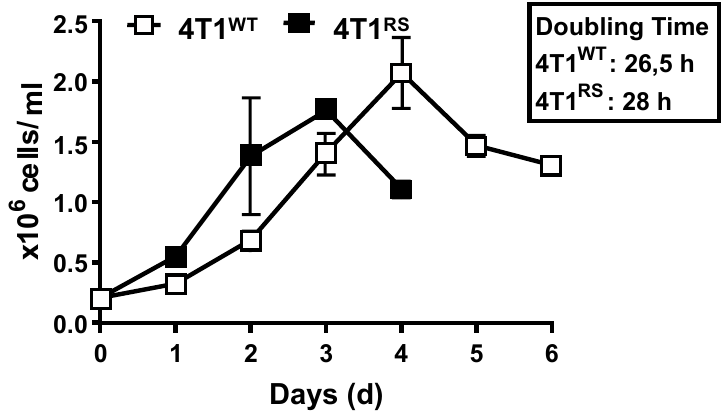

Supplementary Fig. S1: Growth curve of non-irradiated 4T1^WT^ control and hFI-treated 4T1^RS^ cells.

Cell numbers were determined microscopically by cell counting. Data are presented as mean ± SD from two independent experiments, each performed in biological duplicates (n=2, N=2).

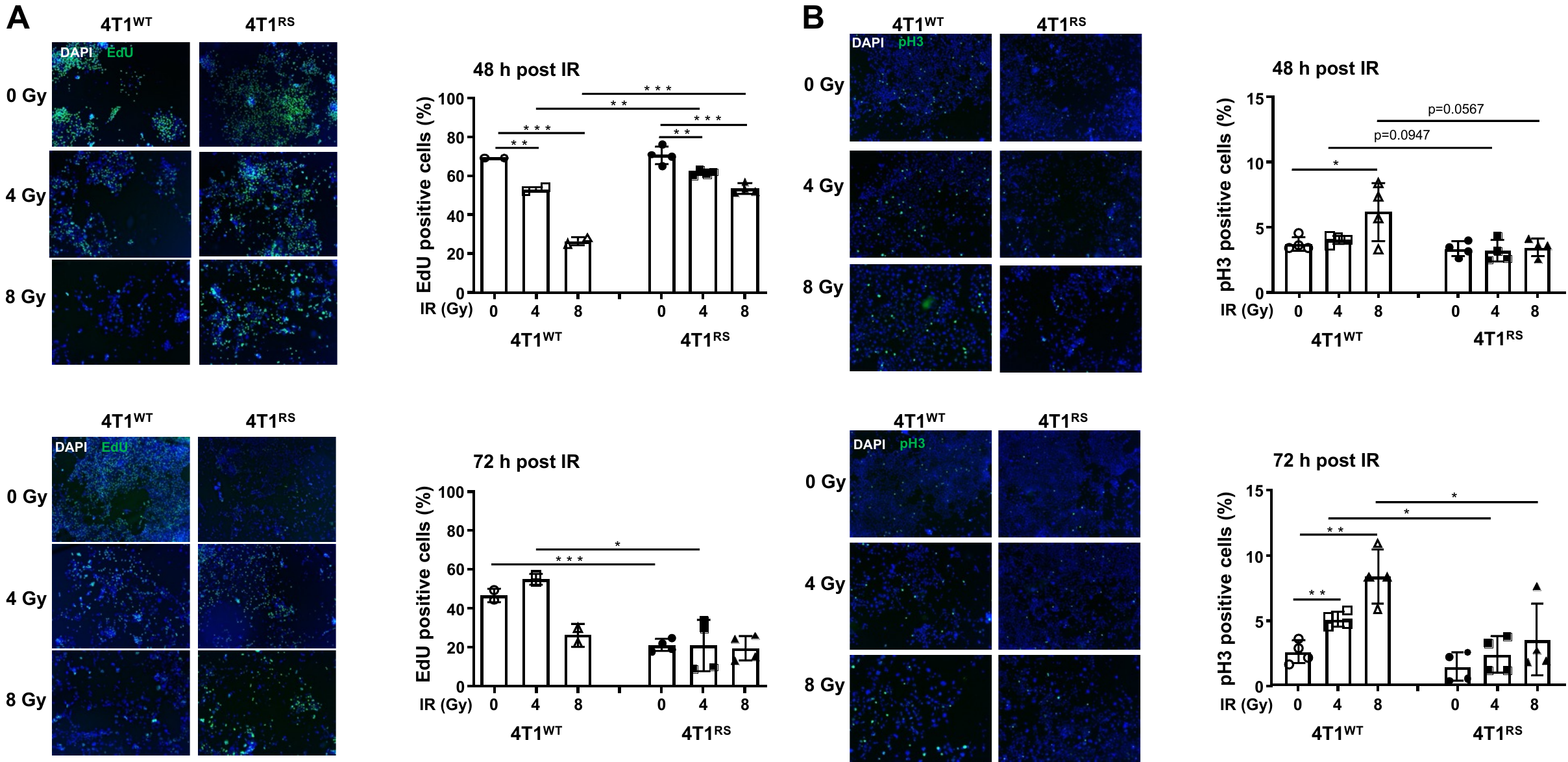

Supplementary Fig. S2: Comparative analyses of proliferation following irradiation treatment in vitro.

**A:** Logarithmically growing 4T1^WT^ and hFI-selected 4T1^RS^ cells were irradiated with 4-8 Gy. After a post-incubation period of 48 h and 72 h cell proliferation was assayed by EdU staining as described in methods. Left panel: Representative pictures. Right panel: Quantitative data are the mean ± SD from n=1-2 independent experiments each performed in biological duplicates (N=2-4). Statistical analysis was performed using unpaired, two-tailed Student´s t-test. p ≤ 0.01; ***p ≤ 0.001 (irradiated cells as compared to the corresponding non-irradiated control).

**B:** Logarithmically growing 4T1^WT^ and hFI-selected 4T1^RS^ cells were irradiated with 4-8 Gy. After a post-incubation period of 48 h and 72 h the mitotic index was determined as described in methods. Left panel: Representative pictures. Right panel: Quantitative data shown are the mean ± SD from two independent experiments each performed in biological duplicates. Statistical analysis was performed using unpaired, two-tailed Student´s t-test. *, p ≤ 0.05; **, p ≤ 0.01 (irradiated cells as compared to the corresponding non-irradiated control).

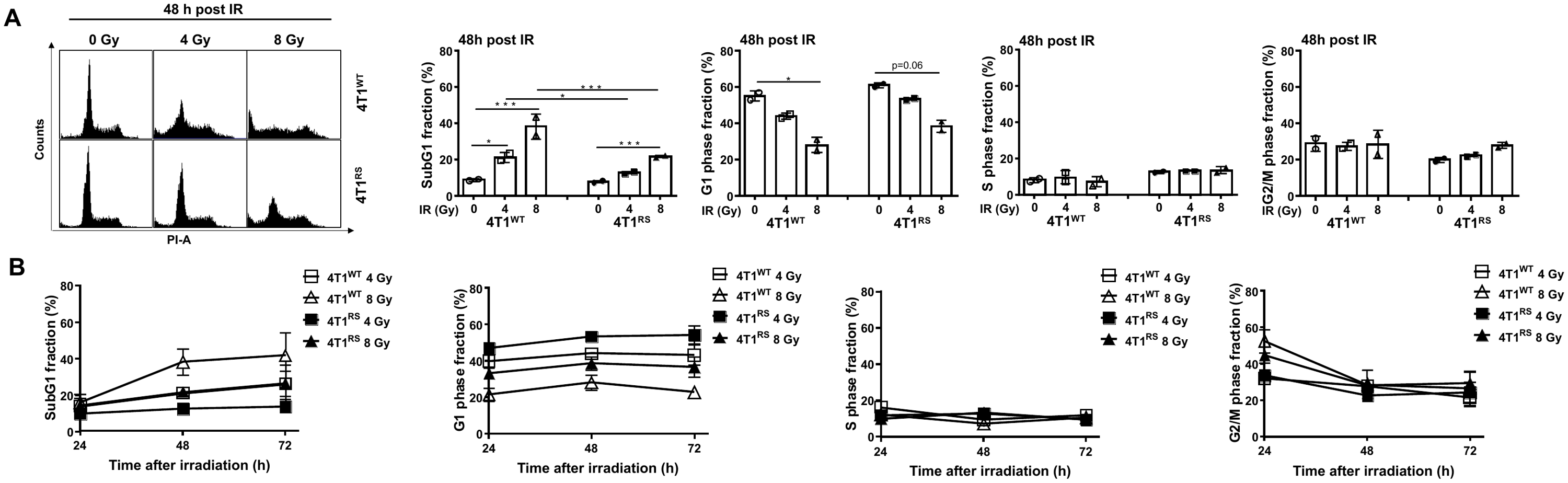

Supplementary Fig. S3: Analysis of alterations in cell cycle distribution following IR treatment.

Logarithmically growing non-irradiated 4T1^WT^ and hFI-treated 4T1^RS^ cells were irradiated with 4-8 Gy. After a post-irradiation time of 48 h, cell cycle distribution was monitored by flow cytometry-based method.

**A:** Left panel: Representative data. Right panel: Quantitative histogram depicting the percentage of cells present in SubG1 fraction, G1-phase, S-phase and G2/M-phase. Data are the mean + SD from two independent experiments each performed in biological duplicates. Statistical analysis was performed using One-way ANOVA . *, p ≤ 0.05; ***, p ≤ 0.001.

**B:** Time kinetic analysis of cell cycle distribution of non-irradiated 4T1^WT^ and hFI-treated 4T1^RS^ cells 24-72 h after irradiation with 4-8 Gy. Data are the mean ± SD from n=2-9 independent experiments each performed in biological duplicates.

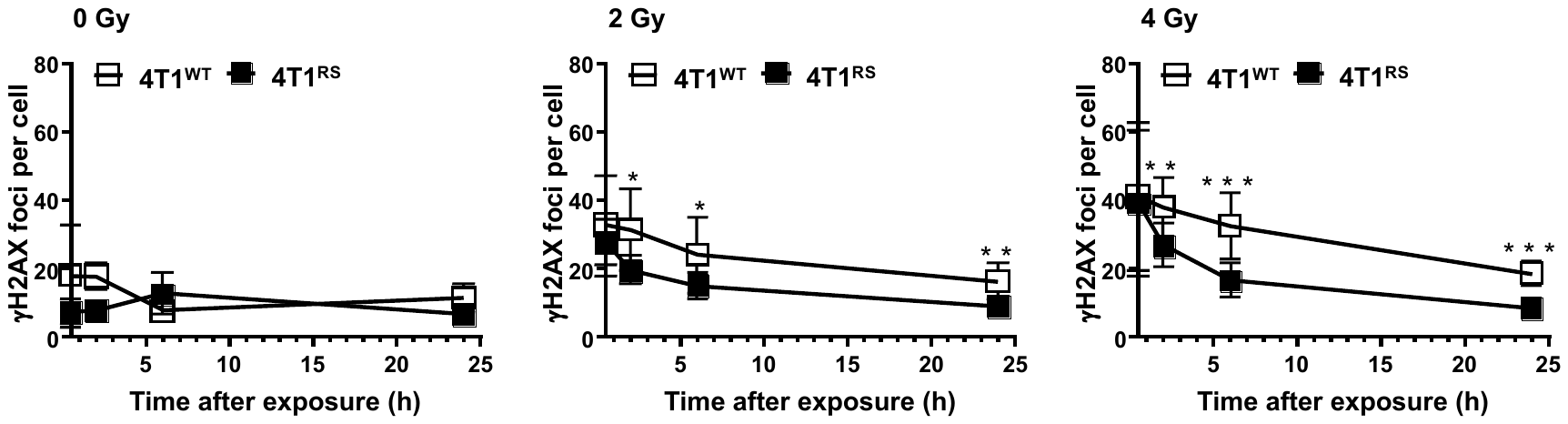

Supplementary Fig. S4: Formation and repair of IR-induced DNA damage.

Logarithmically growing non-irradiated 4T1^WT^ and hFI-selected 4T1^RS^ cells were irradiated with 2-4 Gy. 30 min, 2 h, 6 h, and 24 h after IR treatment the formation of nuclear γH2AX foci was analyzed by immunocytochemistry as described in methods. Number of nuclear γH2AX foci are shown as the mean ± SD from n=4-5 independent experiments each performed in biological duplicates. ≥50 cells were analyzed per experimental condition. *, p ≤ 0.05; **, p ≤ 0.01; ***, p ≤ 0.001.

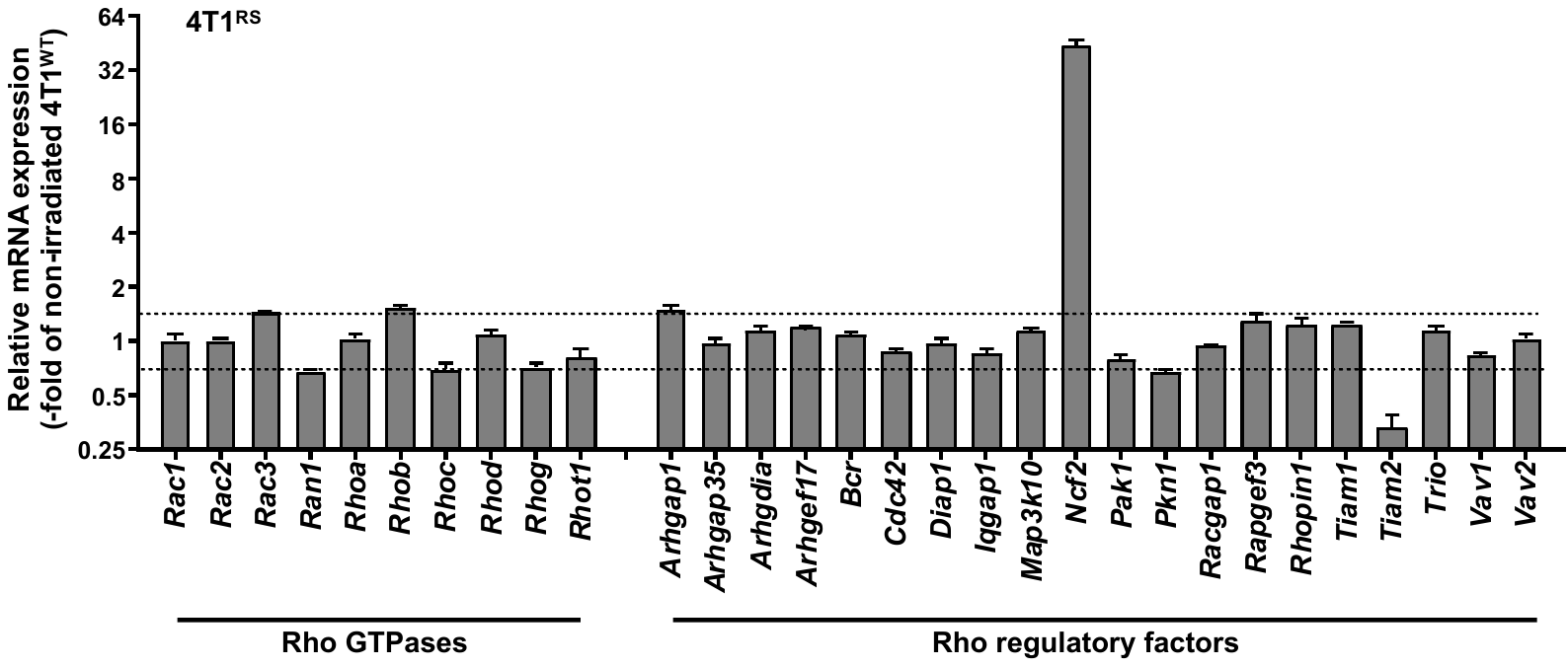

Supplementary Fig. S5: hFI-induced gene expression of Rho GTPases and Rho-regulatory factors.

The mRNA expression of Rho GTPase-related factors (Rho GTPases, Rho-regulatory factors (GEFs, GAPs) was comparatively analyzed in non-irradiated 4T1^WT^ cells versus irradiated 4T1^RS^ cells 24 h after 8 Gy. Relative mRNA expression in radioselected 4T1^RS^ was related to that of non-irradiated 4T1^WT^ cells, which was set to 1.0. Changes in mRNA expression levels of ≥1.5 and ≤0.7 are considered as biologically relevant and marked with a dashed line. Data shown are the mean ± SD from technical duplicates obtained from pooled samples of biological triplicates.

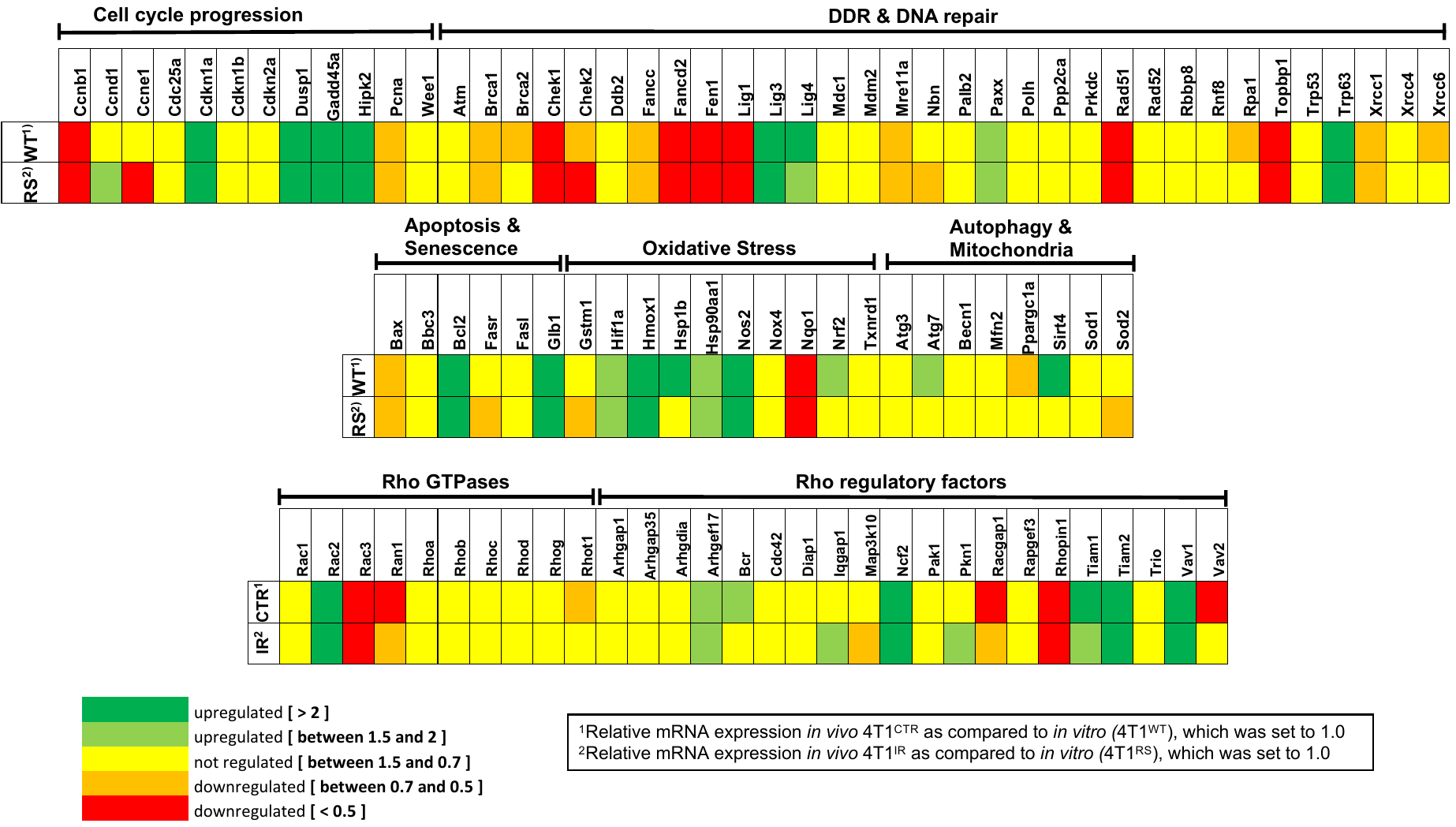

Supplementary Fig. S6: Graphical illustration of alterations in gene expression under in vitro versus in vivo situation in non-irradiated control cells or following hFI.

Basal mRNA expression level of a subset of susceptibility-related genes (involved in the regulation of cell cycle progression, DDR and DNA repair, apoptosis/senescence, oxidative stress response or autophagy/mitochondrial functions) and Rho GTPase related factors (Rho GTPases, Rho-regulatory factors (GEFs, GAPs)) were comparatively analyzed in *in vitro* versus *in vivo* growing 4T1 cells. Relative mRNA expression in *in vivo* growing 4T1^CTR^ and 4T1^IR^ tumors was related to that of *in vitro* growing non-irradiated 4T1^WT^ and hFI-selected 4T1^RS^ cells, respectively, which were set to 1.0.
